# Intramolecular competition generates pulsatory protein activity shaped by light, temperature, and evolution

**DOI:** 10.1101/2025.04.08.647774

**Authors:** Zikang (Dennis) Huang, Malvin Forson, William Benman, Kevin H. Gardner, Lukasz J. Bugaj

## Abstract

Proteins are information processors, but their computations are typically considered at steady state. Here we find that individual proteins can dynamically encode information about their environment and that such response dynamics have been conserved throughout evolution. The fungal protein BcLOV4 exhibits pulsatory light responses shaped by the magnitude of environmental light and temperature. Response adaptation resulted from competitive interactions between domains that sensed either light or temperature. Temperature-sensing was encoded in a modular domain and could be tuned by mutations within co-evolved loops. Photo-thermal response dynamics were conserved in homologues from fungi that diverged >300 million years ago, and the characteristic temperature of pulsatory responses had adapted to match the ecological niche of the hosts, ranging from Antarctica to thermal ponds. These findings uncover a class of dynamic proteins, determine molecular principles of time-varying protein activation, and suggest functional importance for light- and temperature-conditioned protein activity pulses.

**One-Sentence Summary:** Individual proteins can dynamically encode information through interactions between their component domains, revealing principles for complex signal processing in natural and engineered proteins.

## Introduction

Proteins serve as information processors in the cell, integrating combinations of internal and environmental cues to execute an appropriate response (*1*). Both natural and engineered proteins can perform complex processing, for instance acting as logic gates that execute a steady-state response to combinations of inputs (*2*–*4*). In principle, proteins could also process information with non-steady-state computation like dynamic encoding, where input information is transduced into a temporal pattern of an output. However, to date dynamic encoding has generally not been recognized within individual proteins. One exception is voltage-gated ion channels, which rapidly open upon stimulation but subsequently auto-inactivate due to intramolecular domain interactions ^9^. However, it is unclear whether nature has evolved pulsatory activity within proteins more broadly, including with more flexible control of activation patterns and more generalizable output activity compared to ion flow.

Although largely unexplored within proteins, dynamic encoding is commonplace within networks (*5*). Pulse generation is one common form of dynamic encoding, playing a central role in essential sensory behaviors like response to growth factors(*7*), adaptation to stress(*8*–*10*), and recognition of chemical gradients (*11*). Corrupted pulsatory responses of signaling, in turn, have been associated with disease including cancer (*12*). Pulses can be generated by networks that achieve a fast activation phase followed by slower inactivation, implemented by sequential antagonistic interactions. Such dynamics could be encoded in networks of any scale, including in networks of domain interactions within an individual protein.

Here we explore the potential for such regulation within BcLOV4, a photoreceptor from the fungus *Botrytis cinerea*. BcLOV4 generates a pulse of activity lasting tens of minutes in response to constant blue light in heterologous cells ^10,11^, though whether the protein itself is sufficient for pulse generation — or whether additional cellular factors are required — is not understood. What is already clear is that activation dynamics are dependent on a second environmental stimulus: temperature. Activity pulses are observed at elevated temperatures (30-40°C), but below this range adaptation slows and photoactivation is sustained ^11^. Intriguingly, BcLOV4 photostimulation results in its clustering and membrane translocation, both of which are either adaptive or sustained depending on the combined light and temperature inputs ^12,13^. However, we do not understand how clustering and translocation are coordinated, nor how their dynamics respond to multiple environmental inputs.

Using a combination of *in vitro, in cellulo, in silico*, and phylogenetic approaches, we show that BcLOV4 generates activation dynamics as a single protein through competitive intramolecular interactions between sensory domains that respond to either light or temperature. A thermodynamic model based on domain interactions recapitulated this behavior and predicted non-intuitive consequences of mutations and conditions that perturbed the energetics of the underlying protein states. We then uncovered the identity of the relevant domains and their interactions, finding a multimerization region that is essential for membrane binding and that is subject to competing regulation by both a light-sensitive domain (LOV) and a modular temperature-sensitive domain (TSD). We further identified coevolved regions in the TSD that govern the thermal response profile of BcLOV4 dynamics. Finally, eight additional homologues of BcLOV4 retained dynamic response to light and temperature, and the characteristic thermal response of these proteins had shifted to match the ecological niche of the host, implying the functional importance of the dynamic response within this unique class of proteins.

## Results

### A thermodynamic model captures response dynamics of BcLOV4 across light and temperature conditions

BcLOV4 is a 595 amino acid protein from the plant pathogen *Botrytis cinerea* comprising three main globular domains: 1) an N-terminal regulator of G-protein signaling (RGS) domain, 2) a light-sensing light-oxygen-voltage (LOV) domain, and 3) a C-terminal domain of unknown function (DUF) **(Figure 1A)** ^10^. BcLOV4 responds to light by oligomerizing and translocating to the plasma membrane ^6,12^, and translocation is mediated by electrostatic interactions of an amphipathic helix (AH1) located between the LOV and DUF domains **(Figure 1A)**^10,13^. Unique among known photoreceptors, BcLOV4 is also sensitive to temperature, such that the activated protein becomes subsequently inactivated as an increasing function of temperature **(Figure 1B)** ^11^.

**Fig 1.**
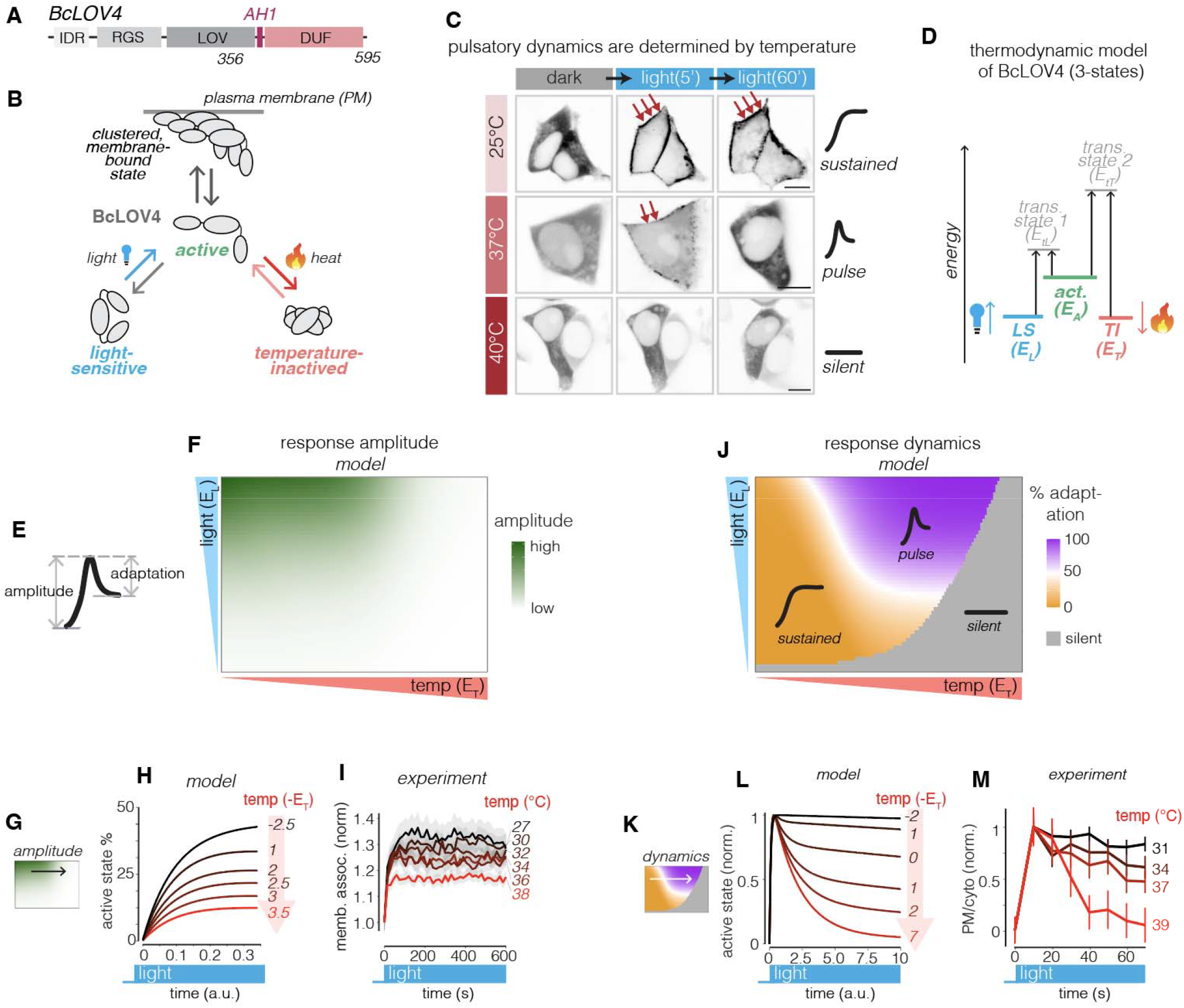
Figure 1: BcLOV4 exhibits 3 classes of dynamic photoresponses that can be captured by a thermodynamic equilibrium between 3 states. (**A**) BcLOV4 domain arrangement. IDR, intrinsically disordered region. RGS, regulatory G-protein subunit. LOV, light-oxygen-voltage. AH1, amphipathic helix 1. DUF, domain of unknown function. (**B**) Hypothesized BcLOV4 state transitions. (**C**) BcLOV4 exhibits 3 classes of response to light and temperature: sustained, pulsatory, or silent. GFP imaging was shown. Also see **Movie S1**. Scale bar = 10 um. Light intensity 23.6 mW/cm^2^, 50% duty cycle with optoPlate stimulation. (**D**) A thermodynamic model describes the three hypothesized states that implement these three response classes. Light stimulation is modeled by increasing energy of the LS state. Temperature increase is modeled by decreasing the energy of TI state. (**E**) Definition of metrics used to quantify dynamic responses. (**F**) Model of how response amplitude changes as a function of light and temperature. (**G**) Testing predictions of response amplitude. Model (**H**) and live cell imaging (**I**) of BcLOV4 amplitude at varying temperatures. Error bars represent SEM of ∼600 cells. (**J**) Model of how response dynamics change as a function of light and temperature. See **Model** section for details. (**K**) Testing predictions of response dynamics. Model (**L**) and experiment (**M**) of BcLOV4 light-response dynamics under different temperatures. In experiments (**I**,**M**), cells were kept under target temperatures for 2 hours using the thermoPlate prior to illumination. Error bars represent SEM of ∼600 cells.

Together, these behaviors implement at least three classes of activity in response to a constant light stimulus **(Figure 1C)**. At low (<∼30°C) temperatures, BcLOV4 activity is sustained. At intermediate temperatures (30-38°C), BcLOV4 becomes pulsatory, rapidly activating but subsequently inactivating over a slower timescale, despite the persistence of blue light. However, at still higher temperatures (>∼40°C), BcLOV4 loses its response to light, which we term a “silent” response. This silent response was not predicted by a prior model of BcLOV4 activation dynamics ^11^, suggesting the need for a new conceptual framework.

To account for these three response classes within a single protein, we hypothesized that BcLOV4 exists in equilibrium between three main states: the light-sensitive (LS) state that can be activated by light; the active state (act.) that allows oligomerization and membrane binding; and the temperature-inactivated (TI) state, which does not respond to light but does respond to temperature changes (**Figure 1B,D**) ^14^. In this framework, the energy states of the LS and TI states are governed by external inputs. Light stimulation raises the energy of the LS state, while temperature regulates the energy level of the TI state, increasing energy level at low temperatures and decreasing at high temperatures.

We implemented this model with ordinary differential equations and observed whether it qualitatively recapitulated BcLOV4 responses to light stimulation. We additionally included a 4th, irreversibly-inactivated (IS) state to account for a previously observed irreversibility that occurs over slow timescales and is a function of light stimulation (**Figure S1**). We modeled the amplitude of activation by quantifying the maximum active state population after light stimulation, and dynamics were quantified by the degree of adaptation, defined as the percentage drop from the max amplitude after the system reached equilibrium **(Figure 1E)**.

Modeling response amplitude showed that 1) at a given temperature, amplitude rose as an increasing function of light, and 2) at a given level of light stimulation, the amplitude of activation *decreased* with increasing temperature, failing to respond at the highest temperatures **(Figure 1F)**. This decreasing and ultimately silent response occurs because the TI state becomes sufficiently favorable to depopulate other states, including the otherwise light-activatible protein in the LS state. Live cell microscopy of stimulation at different temperatures confirmed the decreasing response amplitude as a function of temperature **(Figure 1G-I, S2)**.

Modeling response dynamics revealed a more complicated relationship in how light and temperature shape the temporal response pattern **(Figure 1J)**. At low temperature, BcLOV4 activity is sustained. However, as temperature increases and stabilizes the TI state, the sustained response becomes progressively more pulsatory because transitioning to, and remaining in, the TI state becomes increasingly favorable. Again, live cell microscopy confirmed that, at a given light stimulation level, BcLOV4 membrane translocation shows stronger adaptation with increasing temperature, consistent with prior results ^11^. Our model also captures that adaptation increases as a function of both light and temperature, as shown previously **(Figure 1K-M)** ^11^.

### The equilibrium between states shapes BcLOV4 light response

The thermodynamic model made several testable predictions. First, equilibrium between states suggested that, after transitioning into the TI state, the active state could be repopulated by simultaneously lowering the temperature while maintaining illumination ^14^, where the magnitude of active state repopulation would be a function of light intensity (**Figure 2A,B**). Indeed, live cell microscopy after BcLOV4 adaptation to the TI state at 37°C and subsequent cooling to 25°C showed a reappearance of membrane localization, and localization was strongest under higher blue light intensity **(Figure 2B-C)**.

**Fig 2.**
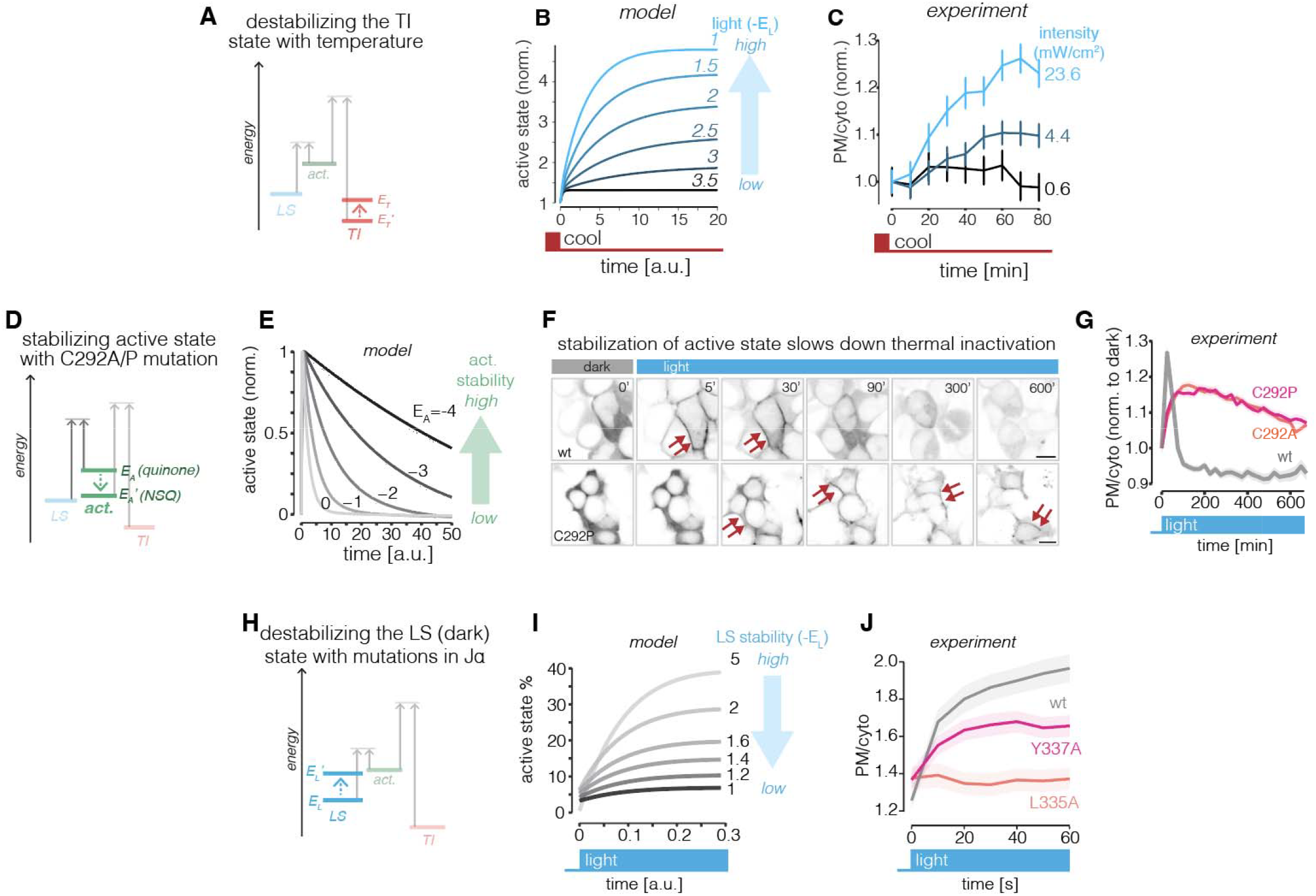
The equilibrium between states shapes BcLOV4 light response. **(A)** Destabilization of the temperature inactivated (TI) state by decreasing temperature. Model (**B**) and live-cell imaging (**C**) of active state population upon decrease in temperature under variable levels of constant light. The cells were pre-illuminated to populate the TI state (see **Methods**) and then cooled to 27°C under different intensities of light stimulation. Error bars represent SEM of ∼300 cells. **(D)** Stabilizing the active state (act.) by mutation of C292 to promote active-state formation of the stable neutral semiquinone (NSQ). **(E)** Simulation of light-activation with a stabilized active state shows slower auto-inactivation. **(F**,**G)** Experimental validation of active-state mutants. BcLOV4(wt) and BcLOV4(C292P or C292A) tagged with mCherry were illuminated at 37C with constant light using the optoPlate LEDs (23.6 mW/cm^2^). Data represent mean +/-SEM of ∼300 cells. **(H**,**I)** Destabilizing the light-sensitive (LS) state is predicted to result in higher basal membrane binding in the dark but decreased maximal binding in the light. () Experimental validation in HEK 293Ts with LS-destabilized variants with point mutations that weaken LOV-Jα interaction. Data represent mean +/-SEM of ∼60 cells. Scale bar = 10 µm.

A second prediction was that stabilization of the active state should decrease the rate of thermal inactivation (**Figure 2D**). We stabilized the active state through mutation of the critical LOV cysteine (C292 in BcLOV4) which forms a covalent adduct with the C4a carbon of a photoexcited flavin mononucleotide (FMN) cofactor ^15^. Natural or engineered LOV domains that lack this critical cysteine can still be photoactivated through alternative photochemistry, likely the formation of a FMN neutral semiquinone (NSQ) that has substantially longer dark-reversion kinetics relative to the canonical quinone form, indicating a more stable form of the active state and/or a higher transition energy barrier^16^. Consistent with this prediction, BcLOV4 variants containing either a C292A or C292P mutation remained membrane localized at high temperatures (37°C), substantially longer than wildtype wt BcLOV4, with strong localization still observed after more than 10 hrs of stimulation (**Figure 2F-G**). Importantly, these C292A/P mutations also resulted in measurable basal (dark) levels of membrane localization, consistent with model predictions of a stabilized active state (**Figure S3**) ^16^. The kinetics of light activation were also slowed for the C292A/P mutants, consistent with both a stabilized energy level and an increased energy barrier for transition to the active state, as previously observed (**Figure 2F-G**)^16,17^.

One final and non-intuitive prediction from our three-state model was the effect of mutations that destabilize the LS state. Studies of a canonical LOV domain, AsLOV2, revealed that photochemical triggering of LOV domains leads to the release of auxiliary alpha-helices (A’α, Jα) from positions on the LOV domain core as part of the activation process ^18,19^. An analogous JL□ helix is seen in secondary structure predictions of BcLOV4 ^10^, suggesting that mutations that weaken the docking of the Jα helix onto the LOV domain core would destabilize the LS state. In a normal 2-state LOV photo-switch, such mutations would result in higher basal levels of activated protein that could be further maximally stimulated with light. However, the BcLOV4 model predicts higher basal levels but a *decreased* maximal stimulation amplitude because the equilibrium of the mutants would push more protein to the light-insensitive TI state even in the dark (**Figure 2H-I**). We tested this prediction with two individual mutations, L335A or Y337A, that alter residues predicted to contribute to interactions between the core LOV domain and Jα helix ^16^. Indeed, both mutants showed slightly elevated basal (dark) membrane localization and *lower* maximal amplitude of light-induced membrane translocation compared to wt BcLOV4, in agreement with model predictions (**Figure 2J, S4**).

Collectively, these series of experiments support the model that BcLOV4 exists in an equilibrium between 3 main states, providing an explanation for how both light and temperature shape its dynamic activation patterns. Notably our results also exclude alternative explanations for adaptation and pulse generation, including oxidative damage of the chromophore or protein degradation, as these explanations would not predict the response dynamics observed above and would not be consistent with the reversibility between states over short timescales.

### Regulated avidity of AH1 is necessary and sufficient for BcLOV4 membrane translocation

To understand the molecular mechanisms that implement the dynamic equilibrium of BcLOV4 states, we first sought to identify the molecular underpinnings of the active state ^10^. BcLOV4 activation is characterized by two functional outputs, membrane translocation and clustering, though whether these two behaviors are related is unclear **(Figure 1B, 3A,B)** ^10,12^. It was initially proposed that light-induced unfolding of the Jα helix revealed an amphipathic helix (AH1) that provided affinity for the membrane ^10^. However, several observations counter this model. First, a short fluorescently tagged AH1 fragment does not visibly localize to the membrane, suggesting that exposure of AH1 itself is insufficient for translocation **(Figure S5A)**. Alternatively, AH1 could contribute to membrane binding by providing weak affinity that is strengthened when combined with additional weak interactions (high avidity). In support of this mechanism, a fusion of AH1 to another weak membrane-binding motif—a polybasic fragment from the STIM1 (referred as STIM1 in the rest of the paper) protein ^20^—localized to the membrane, even though either AH1 or the STIM1 fragment alone was cytosolic **(Figure S5A)**. Furthermore, a fusion of this STIM1 fragment to full-length BcLOV4 localized to the membrane even in the dark, suggesting that dark-state BcLOV4 in the absence of STIM already has a weak propensity for membrane binding **(Figure S5B)**. Importantly, the dark-state membrane association of BcLOV4-STIM1 was lost if positively-charged amino acids in AH1 were removed **(Figure S5B**,**C)**. This result argues that AH1 may in fact be exposed in the dark. Interestingly, the BcLOV4-STIM1 fusion gradually falls off the membrane during light stimulation at 37°C (**Figure S6**), suggesting that although AH1 may not be occluded in the dark, it can be occluded as part of the thermal response that is only licensed after light stimulation. Accordingly, the L335A or Y337A mutations that weaken LOV-Jα undocking and partially mimic light stimulation *reduced* membrane localization of BcLOV4-STIM in the dark, further supporting our 3-state thermodynamic model (**Figure S7**). Collectively, these results suggest that the critical AH1 domain may in fact remain exposed in the LS (dark) state but is caged upon entry to the TI state.

How could light stimulation drive membrane translocation if AH1 is already exposed in the dark? We asked whether light-induced clustering provided a mechanism for enhancing AH1 avidity and resultant translocation. BcLOV4 can cluster in the absence of membrane binding upon mutation of the AH1 region (BcLOVclust ^13^), suggesting that clustering might precede membrane translocation ^12,13^. In support, live-cell imaging of BcLOV4-GFP with sufficiently high magnification and 1 Hz frame rate showed rapid cluster formation in the cytoplasm within seconds of light stimulation, followed by a subsequent loss of clusters and simultaneous increase in accumulation at the membrane (**Figure S8, Movie S2**). To test whether clustering was necessary for translocation, we fused *E. coli* maltose binding protein (MBP), a solubilization tag known to suppress protein condensation and assembly, to the N-terminus of BcLOV4 (**Figure 3C**) ^21^. MBP completely abrogated BcLOV4 translocation at 37°C, but activity was retained at 25°C, suggesting that the fusion was still functional but impaired, indicating a causal role for oligomerization (**Figure 3C-E**). We then tested the sufficiency of clustering by grafting the AH1 region to an orthogonal clustering module, Cry2 (**Figure 3F-G**) ^22^. Light-activated AH1-Cry2 rapidly translocated to the plasma membrane, whereas a mutant lacking two AH1 lysines involved in membrane interactions formed only cytoplasmic clusters (**Figure 3H, I, Movie S3**). Next, we asked whether clustering of BcLOV4 itself was sufficient to drive membrane binding through avidity of a weak membrane-binding domain (**Figure 3J**). We fused the STIM1 polybasic domain to the C-terminus of BcLOVclust. Light stimulation indeed drove the fusion protein to the membrane, whereas BcLOVclust alone formed clusters in the cytoplasm (**Figure 3K, L**) ^13^. Together, our studies argue that light induces conformational changes in BcLOV4 that promotes its clustering, which is both necessary and sufficient for the observed membrane translocation that occurs through increased avidity of weak membrane-binding domains.

**Fig 3.**
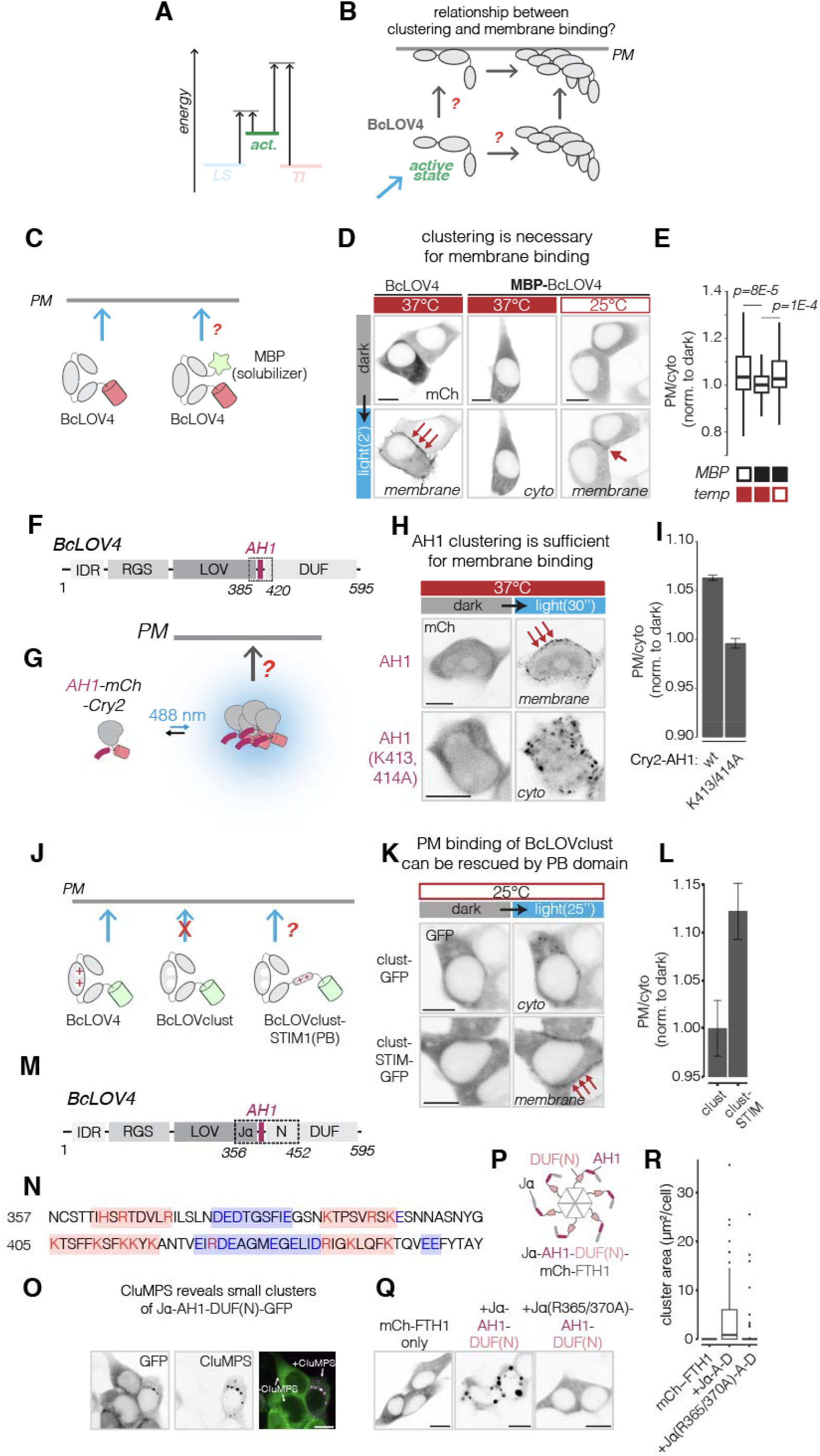
Clustering drives membrane binding of BcLOV4. **(A)** Investigating mechanisms of the BcLOV4 active state. **(B)** Understanding the order and causality between protein clustering and membrane binding. **(C)** Testing necessity of clustering for translocation by fusion of a solubilizing MBP tag that suppresses protein assembly. **(D)** Light-induced membrane binding in HEK 293T cells after 2 min of stimulation. Both constructs were tagged with mCherry. **(E)** Quantification of **(D)**. Data represent mean +/-SEM of ∼70 cells in each group. **(F**,**G)** Testing sufficiency of clustering of the AH1 region for membrane translocation. An extended AH1 fragment (385-420) was fused with Cry2, and protein localization was measured after light-induced clustering. **(H)** Protein localization in HEK 293T cells after 30s of light stimulation. Data show construct from **(G)** and a variant where key membrane-binding lysines were mutated to alanines. **(I)** Quantification of **(H)**. Data represent mean +/-SEM of ∼800 cells. **(J)** Testing whether clustering of BcLOV4 is sufficient to drive weak polybasic domain (STIM1) to the membrane. **(K)** Comparing light-induced localization of BcLOVclust or BcLOVclust-STIM inHEK 293T cells. Both constructs were tagged with GFP. **(L)** Quantification of **(K)**. Data represent mean +/-SEM of ∼130 cells in each group. **(M**,**N)** Patches of charged amino acids are found in the Jα-AH1-DUF(N) region (a.a. 356-452) after LOV domain. Red = positive charge, blue = negative charge. **(O)** Jα-AH1-DUF(N)-GFP appears diffuse in HEK 293T cells but triggers condensation when coexpressed with a CluMPS reporter, indicating the capacity of Jα-AH1-DUF(N) to form submicroscopic clusters. **(P)** Fusion of Jα-AH1-DUF(N) to a 24-meric FTH1 to test the clustering potential of this region. **(Q)** Jα-AH1-DUF(N)-mCh-FTH1 form large condensates in HEK 293T cells, and condensates depend on Jα and two arginine residues therein. Scale bar = 10 µm. **(R)** Quantification of **(Q)** (∼30 cells per group).

### Electrostatic interactions in Jα-AH1 mediate clustering

Our conceptual model implies the existence of a photo-regulatable region that controls clustering. Because LOV domain effectors commonly reside in or downstream of the C-terminal Jα helix, we examined the sequence around this C-terminal region and found patches of charged amino acids (**Figure 3M-N**), a pattern found to promote self-assembly in other protein contexts ^23,24^. We thus asked whether this region, spanning from the Jα helix to the first 20 amino acids on the N-terminus of the Domain of Unknown Function (Jα-AH1-DUF(N)) was sufficient to promote clustering. While a fusion of this region to GFP appeared diffuse, co-expression with a cluster-amplifying CluMPS reporter yielded puncta ^25^, indicating that Jα-AH1-DUF(N) could form submicroscopic assemblies (**Figure 3O**). As further validation, we fused Jα-AH1-DUF(N) to a fluorescently tagged FTH1, a 24-meric subunit of ferritin (**Figure 3P**) ^26^. While FTH1-mCherry alone appeared diffuse, addition of Jα-AH1-DUF(N) resulted in micron-scale puncta, further supporting self-interaction potential of the added region (**Figure 3Q,R**). Substitution of positively-charged arginine residues in Jα with alanines eliminated this clustering potential (**Figure 3Q,R**). Furthermore, mutating 3 lysines in AH1 to negatively charged glutamic acids enhanced clustering, but only in the presence of the Jα, further confirming the dependence of clustering on the charge balance in this region (**Figure S9**). Collectively, these results depict a model where light-induced exposure of the Jα helix reveals critical charged residues that promote clustering and subsequent membrane translocation of BcLOV4.

### The DUF domain mediates temperature sensing

Finally, we sought to understand the molecular nature of the temperature-inactivated (TI) state and how it shapes BcLOV4 response dynamics. We reasoned that pulse generation could occur if the light-sensitive domain (LOV) and a temperature-sensitive domain (TSD) competed for interactions with a shared binding fragment. In the LS state, the LOV core cages a Jα helix, and in the TI state, the adjacent AH1 is caged, though itself is not sensitive to temperature (**Figure S5**). We thus deduced that competitive binding may occur in this Jα-AH1 region, which also contains residues important for oligomerization (**Figure 3M-R**).

To find the temperature sensor, we looked directly downstream of Jα-AH1 in the C-terminal Domain of Unknown Function (DUF) (**Figure 4A-C**). To determine whether DUF contributes to thermo-sensing, we compared membrane localization of the AH1-FTH1 construct in the presence or absence of DUF downstream of AH1. Remarkably, whereas AH1-FTH1 bound the membrane at both high and low temperatures, AH1-DUF-FTH1 bound the membrane at 25°C but not at 37°C (**Figure 4D-F**). Furthermore, addition of the Jα helix upstream of AH1 (Jα-AH1-DUF-FTH1) prevented membrane binding at any temperature, despite the continued avidity provided by FTH1 multimerization, suggesting that Jα plays a strong role in thermal caging (**Figure 4D-F, Movie S4**), in addition to its established role in optical caging. Similar effects of DUF and Jα were obtained with another membrane-binding construct, AH1-DUF-STIM1 (**Figure S10**).

**Fig 4.**
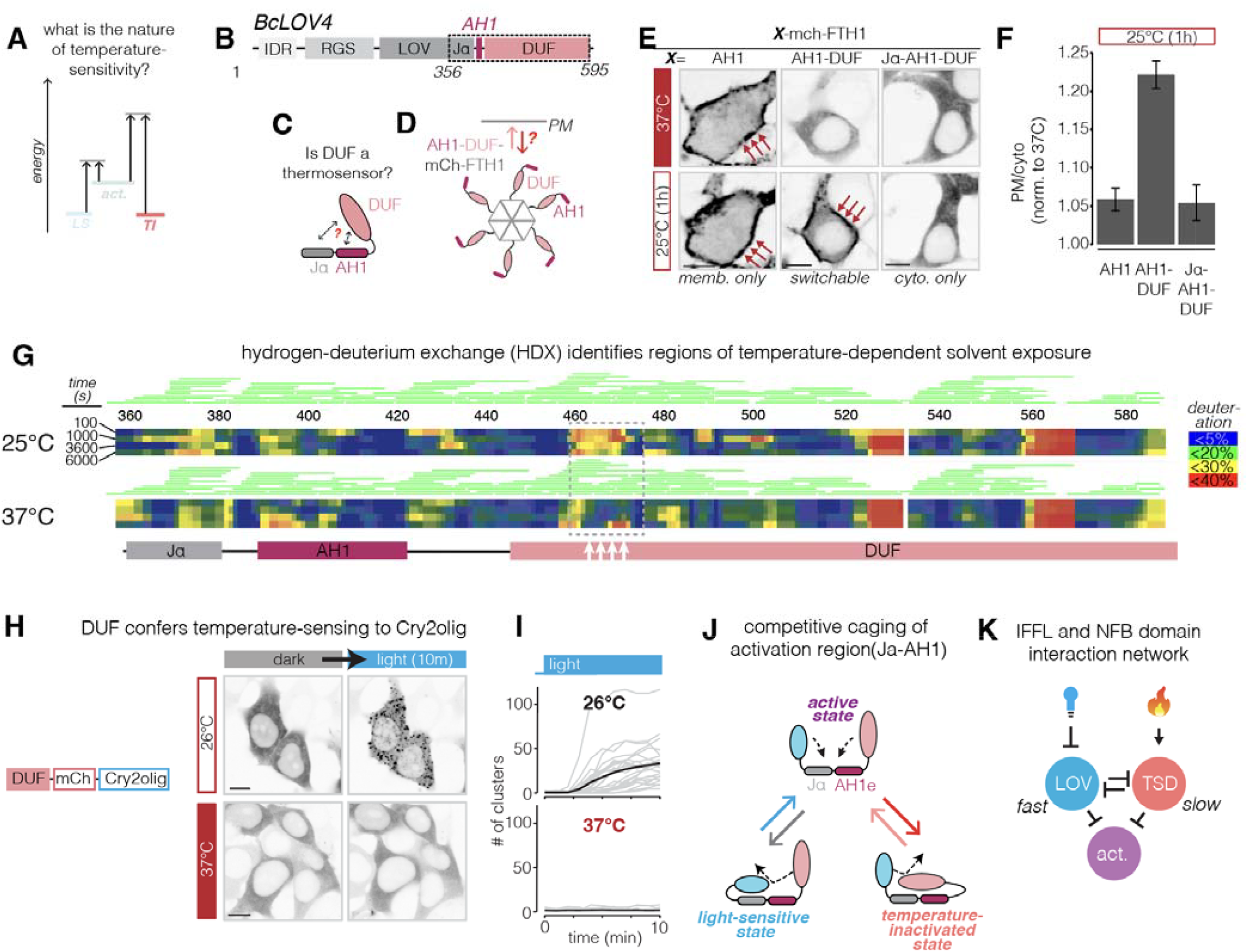
DUF is a modular temperature sensitive domain (TSD). **(A)** Characterizing the mechanisms of the temperature-inactivated (TI) state. **(B**,**C)** The Jα-AH1-DUF fragment (a.a. 356-595) was investigated as a thermosensitive domain. **(D)** Membrane binding of FTH1 fused to Jα-AH1-DUF or truncations thereof was assessed at different temperatures. **(E)** Fusion of FTH1 to AH1 alone (a.a. 85-420) yielded membrane binding at high and low temperatures; addition of the downstream DUF domain (AH1-DUF, a.a. 385-595) resulted in a temperature-switchable membrane binding; further addition of an N-terminal Jα helix (Jα-AH1-DUF, a.a. 356-595) resulted in temperature-insensitive cytoplasmic localization. Also see **Movie S4**. Experiment performed in HEK 293T cells. Scale bar = 10 µm. **(F)** Quantification of **(E)**, showing ratio of membrane localization in the cold state to localization in the hot state. Data represent mean +/-SEM of ∼700 cells. **(G)** Hydrogen deuterium exchange mass spectrometry (HDX-MS) showed temperature dependent conformational changes in the purified Jα-AH1-DUF (a.a. 356-595) fragment. L456-G475 region (highlighted) showed marked temperature induced decrease in deuterium uptake at 37°C compared to 25°C. Time for deuterium exchange is shown on the left. Deuteration level is shown by the heat map. The unique peptides identified by mass spectrometry are indicated by green lines above the heatmap. **(H)** Harnessing DUF for modular thermal control of unrelated proteins (full data see **Figure S19, Movie S7**). A DUF-Cry2olig fusion was tested for clustering propensity under different temperatures (pre-incubation for 2 hours). Scale bar = 10 µm. **(I)** Quantification of **(H)**. Grey traces represent single cells. Black trace represents the mean of all traces. 20-30 cells were quantified for each condition. **(J)** Model for the mechanism of the intramolecular pulse generator, which is implemented by competitive caging of the activation fragment (Jα-AH1) by both the light-sensing and temperature-sensing domains. In the LS state, Jα is caged by the LOV domain, and caging is released by light stimulation. Exposure of this region promotes clustering and subsequent membrane binding but also allows accessibility for thermal caging by the DUF domain, which occurs on slower time scales than light uncaging. (**K**) The domain interaction network of BcLOV4. This architecture generates pulses through a combination of incoherent feed-forward and negative feedback regulation.

To test if temperature sensing is intrinsic to BcLOV4 itself, we looked for temperature-dependent conformational changes in a purified MBP-Jα-AH1-DUF fragment, with MBP added to increase solubility *in vitro*. Limited trypsin proteolysis coupled with mass spectrometry showed differential cleavage patterns in the AH1 and DUF regions when incubated at 25°C vs 37°C. Notably the core of the DUF domain remained stable under both conditions **(Fig S11**,**S12)**. We next subjected the purified fragment to hydrogen-deuterium exchange coupled to mass spectrometry (HDX-MS), which quantifies changes in solvent-accessibility of amino acids by their rate of deuterium incorporation when placed in deuterated water, D_2_O ^27^. While HDX-MS showed no global changes in deuterium uptake performed at 25°C vs 37°C, consistent with the fragment retaining its overall stability, we observed substantial changes in uptake in discrete regions **(Figure 4G)**. This included the 459-472 region which intriguingly showed *decreased* exchange at higher temperature, counter to expectations of *increased* exchange from the mechanism of the exchange reaction ^28^. Few mechanistic explanations can account for such a change, the most probable of which is a temperature-dependent increase in protein interactions in this region. Because of its clear temperature-dependent conformational changes as a purified fragment **(Figure 4G)** and its temperature-dependent functional effects in cells **(Figure 4E)**, we henceforth refer to DUF as the temperature-sensitive domain (TSD).

To assess the modularity of TSD, we asked whether it could endow temperature-sensitivity to unrelated proteins, analogous to how optical control of photosensory domains has enabled modular optogenetic control over numerous and diverse targets ^19^. We thus fused TSD to Cry2_olig_, a variant of Cry2 that clusters robustly at both low and high temperatures. Remarkably, while TSD-Cry2_olig_ clustered at 25°C, clustering was suppressed at 37°C (**Figure 4H, I, S13**). Thus, the temperature-dependent molecular changes in TSD can be harnessed and applied outside the context of BcLOV4 in a modular fashion.

Collectively, these experiments support the model that BcLOV4 pulse generation is implemented through domain competitions that define the three major protein states: In the LS state, Jα is caged by the LOV domain, and caging is released by light stimulation (**Figure 4J**). Exposure of the Jα-AH1 region promotes clustering and subsequent membrane binding but also allows access for caging by TSD, which occurs on slower time scales than the fast light-induced uncaging. This architecture implements a pulse-generating interaction network between the protein domains combining both an incoherent feed-forward loop as well as negative feedback, two network motifs that are commonly found to generate adaptive responses in multi-protein sensory networks **(Figure 4K)**.

### Co-evolution of intra-molecular contacts modulate temperature sensing

We sought to understand how the TSD senses temperature **(Figure 5A)**. Although HDX-MS revealed a temperature-sensitive response in TSD, the responsive region was not within the Jα-AH1 region, implying unexpected intramolecular contacts that mediate temperature sensing. To search for these contacts, we performed a multi-sequence alignment of the corresponding Jα-AH1-TSD region from BcLOV4 homologues and examined the degree of within-protein amino acid covariation using homologues from the same order of Botrytis cinerea, capturing ∼100 million years of evolution ^29,30^ **(Figure 5B)**. Remarkably, the temperature-sensitive regions in the TSD (position ∼460, **Figure 5C**) showed strong co-variation with amino acids near the 560 position **(Figure 5B-D)**. The 460 region contained residues that were both highly conserved and highly variable (**Figure S14A**). The conserved amino acids were crucial to light response, since replacement of any of these conserved residues (Y460, L467, V469) with alanine resulted in proteins that could not respond to light even at low temperatures (**Figure S14B**). By contrast, mutation of the variable residues was not only well tolerated but could also tune the temperature-sensing properties of BcLOV4 (**Figure S14C)**. N466I lowered the temperature at which optogenetic responses could be observed, while N466A and N466M mutations allowed stronger BcLOV4 light responses at both high and low temperatures compared to the wild-type protein (**Figure S14C**). Notably, the I, M, and A substitutions are all found in BcLOV4 homologues, implicating their functionally significant impacts on light- and temperature-sensing.

**Fig 5.**
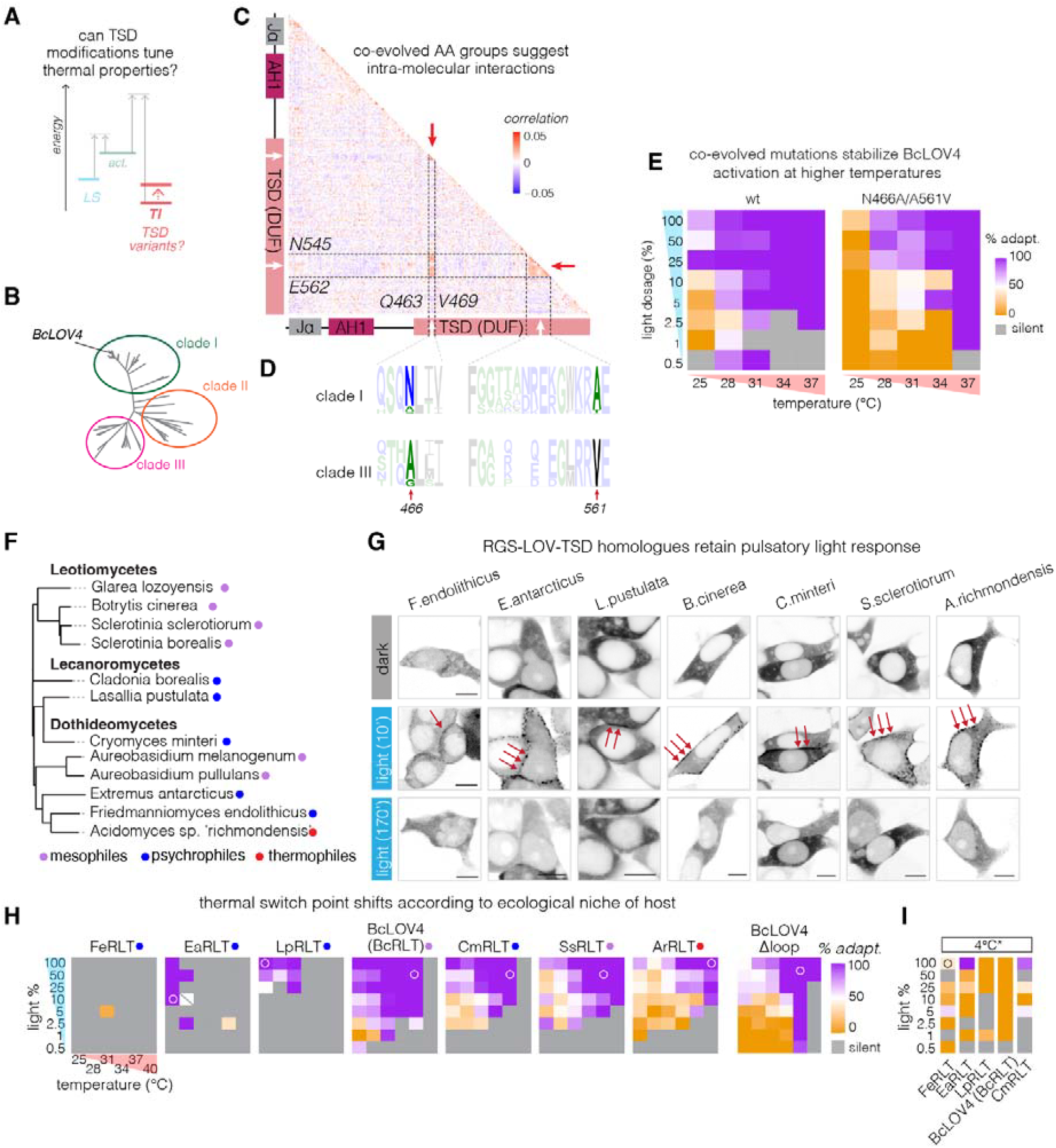
3-class photo-thermal response is evolutionarily conserved and adapted to broad ecological niches. **(A)** Identifying modifications in the temperature-sensitive domain (TSD, previously DUF) that can shift characteristic response temperatures. **(B)** Phylogenetic tree of 69 closely related RGS-LOV-TSD (RLT) homologues. **(C)** Covariation analysis of the homologues in **(B)** revealed fragments of DUF that changed in concert through evolution. These fragments overlap with temperature-sensitive fragments identified through HDX, suggesting functional interactions. **(D)** The indicated amino acids of homologues in clade I (containing BcLOV4) and clade III showed clear covarying patterns. The size of each letter is proportional to the number of instances within the clade. **(E)** Grafting co-varying amino acid pairs (N466A/A561V) onto BcLOV4 shifts the photo/thermal boundary at which responses change from sustained to pulsed. Data represent quantification of pulse shape from live cell imaging of single cells under the indicated light and temperature conditions (see **Figure 1J** and **Methods**). ∼250 live cells were quantified for each grid square. **(F)** 11 homologues of RLT from hosts with known ecological records. The phylogenetic tree was constructed based on the LOV domain. **(G)** 8 RLT proteins (in addition to BcLOV4) showed pulsatory response (see also **Fig S21**), all of which were tagged with GFP. **(H)** Systematic characterization of photo-thermal response under broad light and temperature conditions of 6 variants and BcLOV4 Δloop, which represents a deletion of 544-555 loop region that mimics an ancestral variant before the loop expansion event during evolution. The heatmaps are in the order of increasing switch temperature between the class I (sustained) and class II (pulsatory) response. Ecological temperature of hosts is indicated similarly as **(F)**. Open circles mark conditions used for representative images in **(G)**. Data represent quantification of live cell imaging, as described in **(E). (I)** Testing the light response at 4°C. Cells were illuminated in a refrigerator and fixed before imaging. See **methods** for more details. ∼1000 fixed cells were quantified for each grid. Light (%) represents the percentage of time stimulated with the optoPlate at intensity of 23.4 mW/cm^2^. Scale bar, 10 µm.

To test the functional impact of potential interactions between the 460 and 560 regions, we investigated the co-evolved N466A/A561V mutation pair, which appeared in ∼7% of our homolog dataset and 63% of N466A-containing sequences (**Figure 5D**). Remarkably, this double mutant dramatically increased the switch point temperature at which BcLOV4 response converted from sustained to pulsatory (**Figure 5E**), whereas the individual mutations alone showed smaller or no effects (**Figure S15**). When expressed in HEK 293T cells, the double mutant could be stimulated to reside on the membrane for over 18 hrs, a substantial increase over the wild-type protein, which could only be maintained at the membrane for 1-2 hours under even optimal conditions (**Figure 5E, S1B, S16, Movie S5**) ^11^. Notably, extended membrane localization was achieved only under low levels of light stimulation (2% duty cycle, 12 mw/cm^2^). Higher light levels (20% duty cycle) led to a higher initial magnitude of stimulation but subsequent membrane dissociation after ∼10 hours, in qualitative agreement with the modeled response landscape at intermediate temperature (**Figure 1F**). The double mutant protein also increased the duration and amplitude of an established BcLOV-based optogenetic probe of Ras/Erk signaling in a separate cell line (**Figure S17**) ^11^.

### The 3-class photo-thermal response is evolutionarily conserved and has adapted to a wide range of ecological niches

Finally, we asked whether the environmentally responsive dynamics of BcLOV4 could be found in homologues from evolutionarily distant hosts. We hypothesized that, if these dynamics were functional, they would not only be conserved, but their characteristic switch point temperature would have evolved to match the ecological niche of the host. We focused on homologues with predicted RGS-LOV-TSD (RLT) domain architecture from hosts with known growth conditions or ecological records (**Methods**), ultimately selecting 11 homologues from 3 distinct fungal classes that diverged up to ∼300 million years ago ^31^ (**Figure 5F**). In addition to mesophiles (like *Botrytis cineria*) ^32–36^, we selected hosts that included Antarctic psychrophiles^37–39^ like *Extremus antarcticus* ^40^ and *Friedmanniomyces endolithicus* ^*41*^, and the thermophile *Acidomyces sp. ‘richmondensis’* ^42,43^, which was isolated from a thermal pond in California (**Figure 5F**).

Upon synthesis and expression in HEK 293T cells, 8 of the 11 homologues could translocate to the plasma membrane when stimulated with light. All of these responders showed pulsatory activation, adapting and returning back to the cytoplasm despite constant stimulation over timescales similar to those observed for BcLOV4 (**Figure 5G, S18**). Having confirmed that dynamic light and temperature responsiveness was a widespread feature, we named these proteins to reflect their species and functional domains (e.g., BcLOV = BcRLT)

Individual homologues had distinct relationships between response dynamics and temperature. To best understand these relationships, we performed a comprehensive imaging screen of the light- and temperature-dependent dynamics of six high amplitude responders from across ecological niches. Most homologues could show sustained, pulsatory, or silent response depending on the light intensity and temperature between 25-40°C, similar to BcLOV4 (BcRLT) (**Figure 5G-H**). However, the characteristic transition temperatures between the three classes were shifted within this range. EaRLT and LpRLT were pulsatory below 30°C and silent above this range, whereas CmRLT, SsRLT, and BcLOV4 could be sustained below 30°C and were pulsatory above it. The thermophile ArRLT showed the most dramatic increase in transition temperatures, showing sustained stimulation across temperatures up to 37°C and only achieving fully adaptive pulses at the highest temperature and light exposure.

The psychrophilic FeRLT showed the weakest, mostly silent responses under the conditions tested (**Figure 5H**). We hypothesized this could be because the protein’s transition temperature had adapted more dramatically than other homologues. We thus performed optogenetic stimulation in a refrigerator, and we assessed activation after rapid fixation and imaging ^44^. Remarkably, FeRLT showed strong and sustained activation at this temperature, as did the other psychrophiles in our panel (**Figure 5I**). Overall a clear pattern emerged, where a homologue’s characteristic transition temperatures increased with the environmental temperature of its host. The one exception to this pattern was CmRLT from the Antarctic *Cryomyces minteri*, which had an elevated transition temperature relative to the mesophile BcLOV4. Of interest, prior work found that *C. minteri* is unusually tolerant of high temperatures up to 60°C, consistent with the elevated thermal profile of CmRLT ^37^.

Sequence analysis revealed that homologues outside of the Leotiomycetes lineage (which includes BcLOV4) have partial to complete deletions in both co-evolved functional regions that were identified by HDX-MS in BcLOV4 (**Figure S19**). Comparison of predicted structures using Foldseek identified these as large unstructured loop regions in BcLOV4 ^45^. These loops emerged more than 100 million years ago and expanded over evolutionary time (**Figure S19**) ^30^. Deletion of the 544-555 loop in the BcLOV4 TSD increased its characteristic switching temperature, mimicking a more thermophilic response and suggesting impairment in thermal caging relative to the unmodified protein under equivalent environmental conditions (**Figure 5G, H, S20**). However, temperature sensing was not confined to these loop regions. Homologues from the Dothideomycetes lineage include both the psychrophiles FeRLT and EaRLT, and the thermophile ArRLT, all of which share deletions in the TSD loops (**Figure S19C**). Notably, whereas FeRLT and ArRLT diverged more recently, the psychrophiles FeRLT and EaRLT showed more sequence similarity in the Jα-AH1-TSD region compared to FeRLT and ArRLT, suggesting convergent evolution in adaptation to colder environments (**Figure 5H, I, S21**).

## Discussion

Collectively, our work shows that pulsatory activation of BcLOV4 and related proteins originates from antagonistic intramolecular interactions between its light- and temperature-sensing domains. These interactions implement distinct protein states that result in either sustained, pulsatory, or silent responses to light stimulation, providing a unique example of how individual proteins can dynamically encode information about their environment. Moreover, BcLOV4 transduces these inputs into both its oligomerization and translocation to the membrane, two distinct behaviors that can flexibly regulate molecular biochemistry within the cell ^11,14,46^. Importantly, the dynamic response scheme of BcLOV4 has been conserved and fine-tuned through over 300 million years of evolution, implying a functional role in its hosts.

The light- and temperature-sensing domains generate a pulse of activity due to competitive caging of an “activation” fragment found in the Jα-AH1 region (**Figure 4J, K**). Although initial mechanistic work on BcLOV4 photoactivation proposed that light stimulation exposes the AH1 binding region ^10^, our present work instead suggests that stimulation exposes charged residues in Jα that stimulate higher-order self-assembly, which subsequently causes membrane translocation due to increased avidity of the AH1 region. At the same time, once undocked from LOV, the Jα-AH1 region can be caged by the C-terminal temperature-sensitive domain (TSD, formerly DUF) at a rate that increases with temperature, thereby reducing clustering and membrane binding. Modeling the equilibria of the multiple protein states captured the key dynamic features of BcLOV4, successfully predicted counterintuitive effects of BcLOV4 mutations, and provided a mechanistic explanation for the observed reciprocal regulation of the LS and TI states in response to either light or temperature.

Our model predicted temperature-sensitive auto-inhibition, which we found to be mediated by the TSD. Combining *in vitro* HDX-MS with phylogenetic sequence analysis, we found two co-evolving unstructured regions that underwent conformational changes upon temperature changes, and replacing individual pairs of amino acids with co-evolved pairs in these regions dramatically shifted the thermal response profiles (**Figure 4G, 5B-E**). While our study uncovers how thermal responses are modulated, further work is required to identify the precise molecular contacts that mediate TSD-based auto-inhibition of the activation fragment, as well as other key residues that implement thermosensing. We also identified eight homologues that show similar response dynamics as a function of both light and temperature, but with thermal switch points that have adapted to match the ecological niche of the host. Collectively, these multiple temperature-sensitive protein domains will help uncover fundamental principles by which proteins respond to temperature changes, and their modularity will facilitate the introduction of thermal sensitivity into unrelated proteins (**Figure 5H, I, S21**).

Why do BcLOV4 and other RLT homologues generate activity pulses? In the case of BcLOV4, its host *Botrytis cinerea* is a pathogen of plants including grapes and strawberries ^47^. Perhaps a pulsed vs sustained or silent response allows *Botrytis* to distinguish the seasons, times of day, or levels of sun exposure to ensure optimal infection and fitness ^48^. While a functional role remains speculative, step-to-pulse dynamics are routinely observed across biological networks and have important consequences, for example in mitogenic regulation ^49^, gradient sensing ^7^, or stress adaptation ^2,4,6^.

In these and most known cases, however, pulse generation emerges from the collective action of intermolecular feedback or feedforward networks ^2^. BcLOV4 and its RLT homologues, by contrast, represent a class of individual proteins that use intramolecular interaction networks to transduce environmental inputs into temporal patterns of output activity. This capacity of a protein for dynamic encoding of multimodal information provides a blueprint for understanding and engineering more sophisticated, responsive, and compact biological systems.

## Acknowledgements

The authors thank: Andreas Möglich and Brian Chow for helpful discussion on mechanisms of LOV domain photocycles; Wendell Lim for discussions on protein modularity and computation; Mia Levine for discussions on evolutionary analysis; Yuzhi Gao for help with cell sorting; Rinat Abzalimov for assisting peptide mass spectrometry; and James Siclari for helpful discussions of thermosensing in multiple protein systems. This work was supported by funding from the National Institutes of Health (R35GM138211 for L.J.B., R01 GM106239 for K.H.G.), the National Science Foundation (CAREER 2145699 to L.J.B.), and the Penn Center for Precision Engineering for Health (CPE4H Pilot grant to L.J.B.). Cell sorting was performed on a BD FACSAria Fusion that was obtained through NIH S10 1S10OD026986.

## Contributions

Z.H., K.H.G, and L.J.B. conceived the study. Z.H. proposed the thermodynamic model and performed all the live cell imaging and simulations. M.F. purified the BcLOV4 fragment and performed and analyzed limited proteolysis and HDX-MS. W.B. characterized membrane association of BcLOV4-STIM1. L.J.B. and K.H.G supervised the work. Z.H. and L.J.B. wrote the manuscript and made figures, with editing from all authors.

## Methods and materials

### Plasmid constructs and molecular cloning

BcLOV4 and SOScat genes were obtained from previous work ^11^. FTH1 was sourced from pCMV-SPORT6_FTH1, obtained from the High-Throughput Screening (HTS) Core at Penn Medicine. Cry2 (only the PHR domain was used in this study) sequence was obtained from previous work ^22^. The point mutations were introduced by using primers with altered sequences. DNA templates were amplified using Q5 hot start polymerase from NEB (M0493S), followed by HiFi assembly (NEB E2621S) with backbone DNA digested with proper restriction enzymes. The assembled products were transformed into chemically competent cells (NEB Turbo, C2984H) and subject to antibiotic selection. The homologue sequences were synthesized as DNA fragments by Twist, followed by the similar cloning process above. All DNA sequences used in the paper are included in the **supplementary material**.

### Cell culture, transfection, viral packaging, and transduction

The LentiX-HEK 293T (TakaraBio, #632180) cells were cultured in DMEM with 10% fetal bovine serum (FBS) and 1% penicillin/streptomycin (P/S), while mouse fibroblast NIH 3T3 cells were cultured in DMEM with 10% calf serum (CS) and 1% penicillin/streptomycin (P/S). The cell culture incubators were set to 37C and 5% CO2 without further specification. Before optogenetic stimulation of Ras-Erk pathway in **Figure S17**, cells were serum starved overnight (5X 80% media change). The constructs were characterized in HEK293 cells using transient transfection without further specifications. The transfection was performed with calcium-phosphate precipitation. For long-term stimulation of BcLOV4 (**Figure 4K**) and optogenetic control of Ras-Erk signaling in 3T3 (**Figure S17**), cells were transduced with corresponding virus prepared by collecting the media from HEK cells 2 days after cotransfection of target plasmid, pCMVdR8.91, and pMD2.G (Addgene # 12259).The positive cells were sorted after expansion (AriaFusion).

### Live cell microscopy and high-throughput profiling of light and temperature response

Cells were seeded into Cellvis 96 well plates (#P96-1.5P) coated with fibronectin (MilliporeSigma™ # FC010 diluted 100x in PBS) 24 hr (another 24 hr with transfection and expression) before imaging. Upon seeding, the cells were spun down onto the bottom of plates with 20g and 20 seconds. Live-cell imaging was then performed using a Nikon Ti2-E microscope with a Yokagawa CSU-W1 spinning disk, 405/488/561/640 nm laser lines, an sCMOS camera (Photometrics), and an environmental chamber with CO2 set to 5% and temperature set to 37C without further specifications (Okolabs). To characterize the temperature sensitivity of BcLOV4 variants, we used thremoplate to control the temperature in 96-well plates while imaging ^6^. The environmental chamber was set to 25C during thermoplate control. The thermoplate was allowed to warm up to the target temperature before inserting into the imaging plates. For characterizing light response under different duty cycles and temporal patterns, optoplate was used to deliver blue light from the top of the imaging plate and the confocal microscope was used for imaging from the bottom ^44^. For imaging with thermoplate and optoplate, 150 ul of culture media was added to each well and 100 ul PBS was added to the gaps of wells to prevent evaporation. For the high-throughput characterization of light response in Figure 5, every other row was used in 96 well plates (#P96-1.5P CellVis) to prevent overheating from optoplate. Experiment at 4°C was carried out in a fridge with optoplate followed by fixing with 4% PFA for 10 minutes and 2 times of PBS washes. The cells were cultured in media with HEPES buffering. They are exposed to light for different amounts of time before fixation to capture a pseudo time course. The fixed cells were imaged with the same settings on the confocal microscope as the rest of the live-cell experiments.

### Immuno-fluorescence staining

For characterizing Ras-Erk signaling, we used Immuno-fluorescence staining of ppERK to characterize the result of optogenetic stimulation. Since BcLOV4 is temperature sensitive, and optoplate stimulation can generate heat, we used the thermoplate to maintain the temperature of each well at 37C (with incubator ambient temperature set to 32C) ^6,11,44^. The time course of signaling induction was obtained by starting the stimulation in different wells at different time points but with the same ending time (fixation). Upon completion of the experiment, the optoplate and the thermoplate were disassembled and cells were immediately fixed with 4%PFA (diluted from 16%) for 10 min in dark. Afterwards, the PFA-containing medium was aspirated and cells were permeabilized with 100 ul 0.5% Triton X-100 for 10 min (Sigma), followed by 100 ul ice-cold 100% methanol at -20°C for 10 min. Then, cells were blocked for 30 min at room temperature in Odyssey Blocking Buffer (LI-COR). Anti-ppErk (CST #4370, 1:400) primary antibody were then diluted in fresh blocking buffer. Blocking buffer was removed and cells were incubated in 50 ul primary antibody solutions for 2 hours. Cells were washed 5X with PBS with 0.1% Tween-20 (Sigma). Then the cells were incubated in secondary antibody (CST #SA5-10034, 1:500) together with DAPI (Molecular Probes, #D1306, 300 nM) for another hour followed by 5X wash with PBS with 0.1% Tween-20 (Sigma) ^44^. Then, the samples were imaged using a Nikon Ti2E epifluorescence microscope with DAPI/FITC/Texas Red/Cy5 filter cubes, a SOLA SEII 365 LED light source.

### Protein purification

The Jα-AH1-DUF region of BcLOV4 (I356-F593) was amplified by PCR and cloned into a pMBP-6His-Parallel bacterial expression vector ^50^. The genetic construct was transformed into competent *E. coli* DH5a cells. All sequences were verified by Sanger sequencing. Recombinant proteins were expressed in *E. coli* BL21(DE3) cells (Stratagene). Cells were shaken (180 rpm) post-induction for 18–22 h at 18°C. Cells were harvested and resuspended in buffer containing 50 mM Tris [pH 8], 500 mM NaCl, 0.5% Triton X-100, 0.5 mM DTT, and lysed by sonication. Lysates were centrifuged at 34,571 xg, 4°C for 45 min. Supernatants were filtered through a 0.22 μm Corning syringe filter and bound to a 5 ml Ni^2+^ Sepharose affinity column (Cytiva). The MBP-6His tagged protein was washed with 10 column volumes of protein wash buffer (50 mM Tris [pH 8], 500 mM NaCl, 0.5 mM DTT, 10% glycerol, 50 mM imidazole) and eluted with protein wash buffer supplemented with 450 mM imidazole. Eluted proteins were further purified by size exclusion chromatography on a Superdex 200 16/40 (Cytiva) with SEC buffer containing 50 mM NaPi [pH 8], 150 mM NaCl, 3% glycerol, 0.5 mM DTT. Protein was concentrated using Amicron centrifugal concentrator with 50 kDa MWCO. Concentrations were determined from the theoretical absorption coefficient, ε_280_= 90760 M^-1^ cm^-1^ for MBP-6His-Jα-AH1-DUF, calculated from the sequence using the ExPASy ProtParam server ^51^.

### Limited proteolysis

Limited proteolysis reactions were performed with 50 μM MBP-6His-Jα-AH1-DUF in a buffer containing 50 mM NaPi [pH 8],150 mM NaCl, 3% glycerol, 0.5 mM DTT. Samples were equilibrated at 25°C and 37°C conditions pre-trypsinolysis. Trypsin was added in a 1:500 v/v ratio to MBP-6His-Jα-AH1-DUF at 25°C and 37°C. 40 μL aliquots of the reactions were quenched with 1 μL of PMSF quench solution (final concentration of 2 mM) at timepoints of 100, 500, 1000, 2000, 3600, 5000, 6000, and 7200 s.

Quenched samples were subjected to SDS-PAGE analysis and visualized using Coomassie Blue stain. 3 μL of each terminated reaction was also analyzed for peptide population composition using HPLC-ESI MS/MS LC-MS/MS. The peptides were resolved using a C18 analytical column (Acclaim 300 C18, 3 μm, 2.1 × 1500 mm, Thermo Fisher Scientific) and analyzed on a Bruker maXis-II ETD ESI-QqTOF mass spectrometer. The resulting MS/MS fragments were analyzed using Bruker COMPASS DataAnalysis by deconvoluting the spectrum for specific peptide m/z signals. Peptide sequences were analyzed and identified by Peptide Analysis Worksheet Freeware Edition (PAWs, ProteoMetrics– freeware edition) and the ExPASy ProtParam server. Predicted masses were compared with experimentally generated masses to corroborate SDS-PAGE results.

### HDX-MS

MBP-6His-Jα-AH1-DUF was purified as described above. The stock samples were concentrated to 40 μM and then microcentrifuged. The stock was equilibrated at 25°C and 37°C for 5 min. Deuterium exchange was initiated by adding 5 μL of protein sample to 70 μL of D_2_O HDX buffer (50 mM NaPi [pD 7.6], 150 mM NaCl, 0.5 mM DTT, 3% glycerol, 99.9% D_2_O) and allowed to proceed for 100, 1000, 3600 and 6000 s at 25°C and 37°C. At the end of each reaction time point, the reaction mix was quenched with 75 μL ice-cold quench buffer (3 M GdHCl, 3% acetonitrile, 0.8% formic acid) and kept on ice through all subsequent steps. Quenched protein samples were immediately injected over an Enzymate BEH Pepsin Column (Waters) to digest the protein, while also desalting the resulting peptides with a C18 Hypersil Gold 3 μm 10×1 mm trap column (Thermo Fisher Scientific). After 3 min post-injection, the peptides were resolved through a C18 analytical column (Hypersil Gold, 50 mm length,1 mm diameter, 1.9 μm particle size, Thermo Fisher Scientific) and injected into a Bruker maXis-II ETD ESI-QqTOF high-resolution mass spectrometer. All resulting data files were then imported into version 3.3 of the HDExaminer software (Trajan Automation) to calculate deuterium uptake profiles.

Prior to starting HDX experiments, “unlabeled” samples were run under identical conditions as HDX except the buffer was H_2_O based to provide masses and retention times of undeuterated peptides as reference. The unlabeled run was performed with MS/MS fragmentation to identify peptides using Bruker COMPASS DataAnalysis 5.3 and BioTools 3.2 software. Every matched m/z and retention time pair was assigned to only one peptide sequence based on accurate mass and MS/MS measurements. This resulted in coverage of about 98% of the protein sequence. All peptide matches were manually confirmed after automatic assignment by HDExaminer. All HDX-MS experiments were conducted in duplicates.

### Image analysis

The membrane binding was quantified using the ratio of the mean of membrane fluorescence and the mean of whole cell fluorescence. The cells were segmented with MorpholibJ, a plugin of imageJ (FIJI). The edge pixels of the segmented objects were defined as membranes. The measurements of pixel intensities were done with CellProfiler scripts. Then the output csv files were imported into customized R scripts with post-processing. For quantifications involving the comparison of dark state membrane association, the raw values are shown (e.g. **Figure 2K**), while for comparing dynamic range of light response, the values normalized to dark state (the frame before light stimulation) are shown (e.g. **Figure 2H**).

The number and area of protein clusters within cells were quantified using a customized Matlab script ^25^. The algorithm first detects the center of clusters by the contrast with the surrounding area and then expands to the boundary of the clusters. The two parameters for center and edge detections were kept the same within the same experiment for comparisons between groups.

The ppERK staining quantification, the cells were segmented with DAPI (nucleus) and then a ring of 5 pixels surrounding the nucleus was taken for intensity measurements in both mCherry (expression of construct) and cy5 (ppERK staining) channels using CellProfiler. Then the output csv files were imported into customized R scripts with post-processing.

### Multi-sequence alignment, co-evolution, and phylogenetic analysis

To search for BcLOV4 homologous that have altered temperature sensitivities, we blasted the sequence (BcLOV4 353-595) using the “Non-redundant protein sequences (nr)” database and took the top 69 hits (**supplemental material**). Then, multi-sequence alignment was performed using clustal-omega and visualized using Jalview. The phylogenetic tree of these sequences were generated using iTOL (**Figure 5B,F**) ^52^. The sequence logo was generated using WebLogo 3 with probability as the unit of y-axis (**Figure 5D**) ^53^. The same sequence (BcLOV4 353-595) was input to Amoai for calculating the co-variation between every other amino acid using default settings (**Figure 5C**) ^29^.The full length RGS-LOV-TSD homologues were blasted with “ClusteredNR” database on NCBI for more divergent hosts (**Figure 5F**). FoldSeek was used for aligning TSD structural homologues (**Figure S19D**) ^45^. Timetree was used to source the divergence time of taxons, which is further verified by the references cited.

### Dynamic modeling

We modeled the state transitions of BcLOV4 with ordinary differential equations (ODEs). The kinetic model is illustrated in **Figure S1**. We used the following equations to model state transitions of BcLOV4:

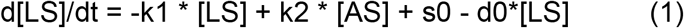

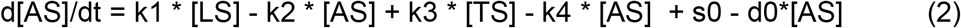

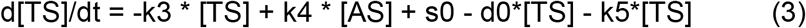

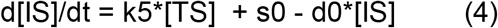

[XX] refers to the concentration of XX state. k1-k5 are the rate constants of state transitions. s0, rate constant for protein synthesis. d0, rate constant for protein degradation.

To simulate the dynamics of the system upon temperature or light changes, initial distribution of the states are acquired through evolving a random distribution towards its equilibrium (or infinite time in ODEs) (**Figure 2B**). Then, the changes on light (k1 is increased to simulate illumination) or temperature (k3 is increased to simulate cooling) are imposed on the initial distribution and induce the transition between states. The energy terms are derived from the correlation between rate constants and activation energy given by:

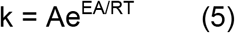

E_A_ can be calculated as the difference between transition states and stable states (LS, AS, TI). Since the energy terms are relative, we set E_act_ = 0 unless specified and calculated other energy terms accordingly from the set of rate constants used in ODEs (**Figure 1H, J**). We also assume the transition states are kept constant under different environmental conditions. Please refer to supplementary file 1 to see the analytical solutions of population distribution at equilibrium, the discussion on how light affects k5, and parameterization details.

