## Supplemental information for "Intramolecular competition generates pulsatory protein activity shaped by light, temperature, and evolution"

**Supplementary Figures**

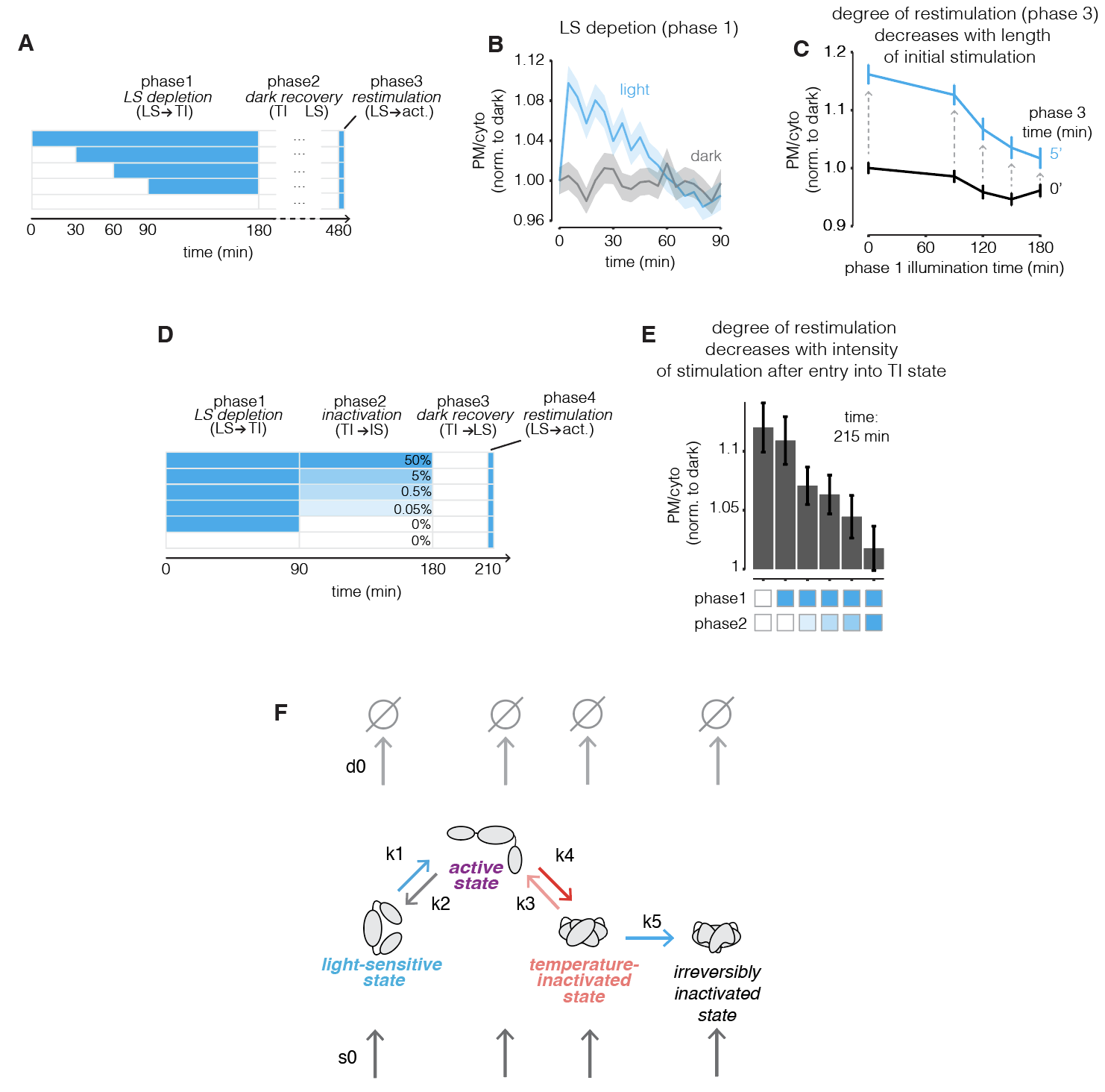
**Fig S1. Kinetic model for BcLOV4 state transitions.** **(A)** Protocol for testing the impact of length of light stimulation on the loss of reversibility of BcLOV4. In phase 1, HEK 293T cells (same for all panels in **Fig S1**) transfected with BcLOV4-mCh were stimulated with 0, 90, 120, 150, 180 min of light at 50% duty cycle (12 mW/cm^2^, same for all panels in **Fig S1**), with light administration using an optoPlate. In phase 2, cells were incubated in the dark for 5 hours. In phase 3, cells were restimulated for 5 min to visualize the restored light-sensitive population. **(B)** Representative traces of membrane localization during phase 1 in **(A)**. BcLOV4 translocated to PM at the beginning of light stimulation and gradually dissociated from PM until its completion around 90 min. The data represent mean +/- SEM of ~550 cells. **(C)** The reversibility of BcLOV4 light response decreases with length of stimulation in phase 1. The black line indicates PM binding after phase 1 as a function of illumination time, while the blue line indicates PM binding after phase 3. The difference between blue and black lines highlighted with gray arrows indicate the reversibility of light response. Data represent mean +/- SEM of ~300 cells on the blue line, ~680 cells on the black line. **(D)** Protocol for testing the impact of the light intensity on the loss of reversibility of BcLOV4. In phase 1, cells transfected with BcLOV4-mCh were stimulated with 50% duty cycle for 90 min to allow for full transition of the LS state to the TI state (as shown in **B**) or were kept in the dark. In phase 2, cells were stimulated for 90 additional mins using different duty cycles as indicated. In phase 3, the cells were incubated in the dark for 30 min. In phase 4, cells were restimulated for 5 min to visualize the degree of restoration of the LS population. **(E)** The PM binding after phase 4 restimulation between different stimulation patterns was compared. The duty cycles used in phase 1 and 2 are indicated by shading under the bars. Data represent mean +/- SEM of ~300 cells. **(F)** k1-k5 are rate constants for state transitions (k1 and k5 are functions of light, k3 is a function of temperature), s0 and d0 are rate constants for protein synthesis and degradation (same for each state for simplicity).

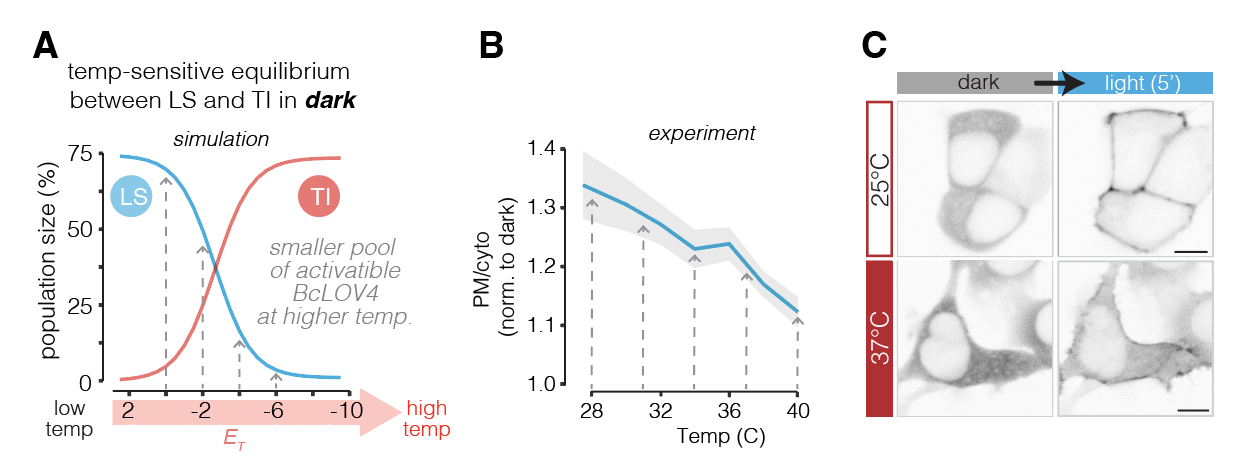
**Fig S2. The effect of temperature on population distribution between states before light stimulation.** (**A**) Light stimulation was simulated by increasing k1 after equilibrating populations under different temperatures (E_T_). Simulation shows a lower amplitude of activation at higher temperatures. (**B**) Cells were incubated under different temperatures for two hours and then stimulated with blue light. The 210s time point was chosen to generate **Fig S2C**. Data show mean +/- SEM of ~530 cells. (**C**) Representative images of (**B**). Scale bar, 10 um.

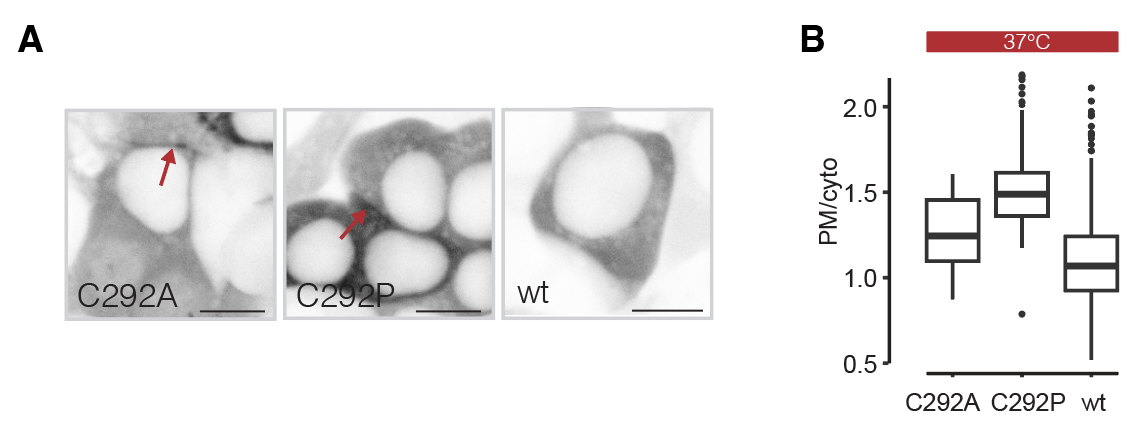
**Fig S3. Mutations of C292 stabilize the active state and result in elevated membrane binding in the dark. (A)** HEK 293T cells were transfected with the indicated BcLOV4-mCh variants and were imaged after incubation at 37°C. Scale bar = 10 µm. **(B)** Quantification of (A). ~130 cells are quantified for each group.

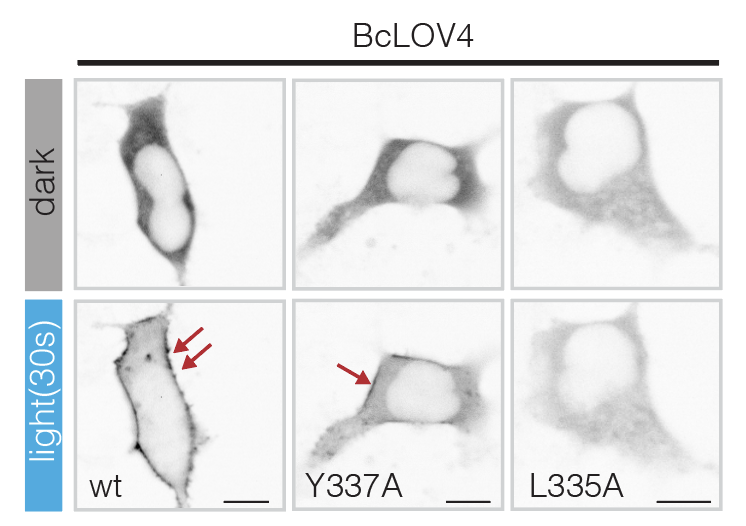

**Fig S4. Destabilization of light-sensitive state results in a decrease in maximal activation.** Representative images for **Fig 2K**. Scale bar = 10 µm.

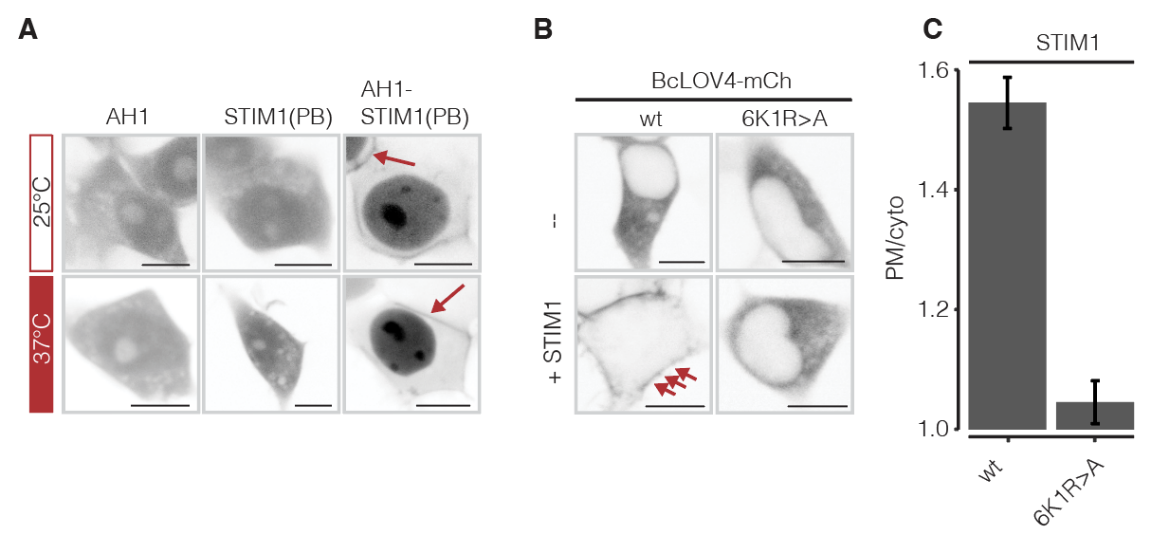

**Fig S5. Evidence that AH1 is exposed in the dark. (A)** AH1 binds the plasma membrane when fused with another weak membrane-binding domain from STIM1. **(B)** BcLOV4 binds PM when fused to STIM1 in the dark in an AH1-dependent manner. **Mutation of 6 lysines and 1 arginine in AH1** (6K1R, BcLOVclust) abrogates membrane binding, suggesting that these amino acids contribute to membrane binding in the dark and are thus exposed. **(C)** Quantification of **(B)**. 6K1R>A refers to mutation of K388, 395, 405, 410, 413, 414A, R393A, which are the positive charges important for PM binding. Data represent +/- SEM of ~70 cells. Scale bar = 10 µm.

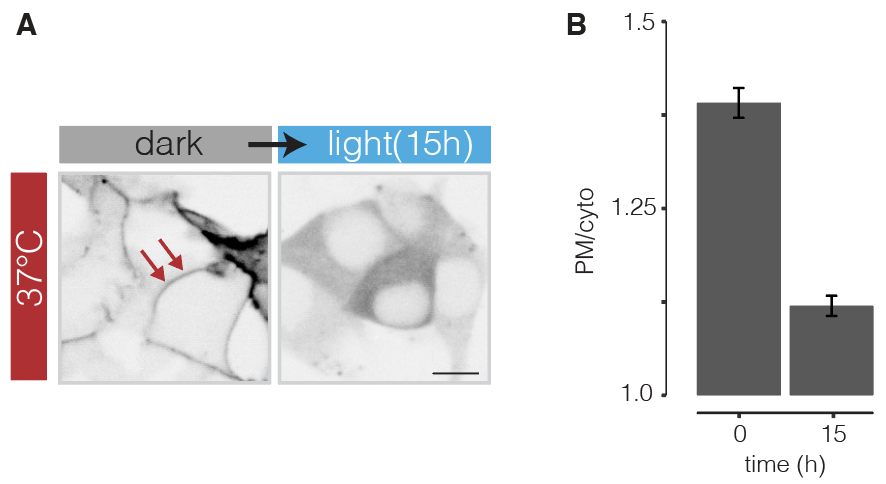

**Fig S6. AH1 is occluded after long term light stimulation at 37°C.** HEK 293T cells transfected with BcLOV4-STIM1 were cultured at 37°C and illuminated for 15 hrs using the optoPlate (5.9 mW/cm^2^ and 50% duty cycle). Data represent mean +/- SEM of ~450 cells. Scale bar = 10 µm.

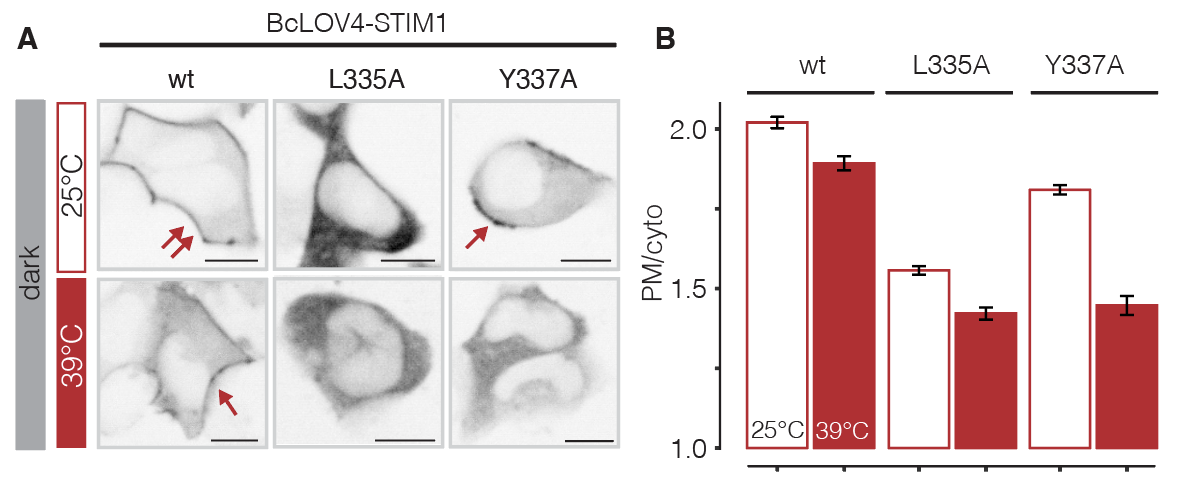

**Fig S7. Destabilization of light-sensitive (LS) state reduces membrane-binding of BcLOV4-STIM1 in the dark. (A)** Either L335A or Y337A weakening LOV core and Jα interactions was introduced to BcLOV4-STIM1. HEK 293T cells transfected with the indicated constructs were incubated at 25°C or 39°C for 2 hours before imaging. **(B)** Data represent mean +/- SEM of ~800 cells (25°C) and ~400 cells for (39°C). Scale bar = 10 µm.

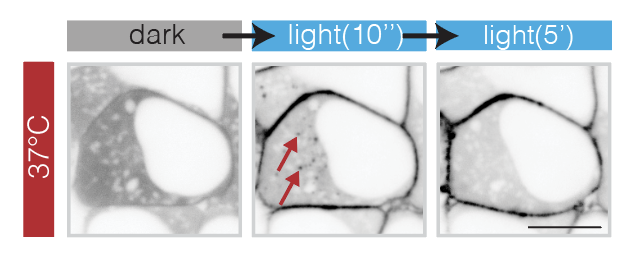

**Fig S8. BcLOV4 transiently forms visible clusters in the cytoplasm within seconds of light stimulation.** High-magnification confocal microscopy reveals the presence of transient clusters, which disappear likely due to translocation to the membrane. Scale bar = 10 µm. See also **Movie S2**.

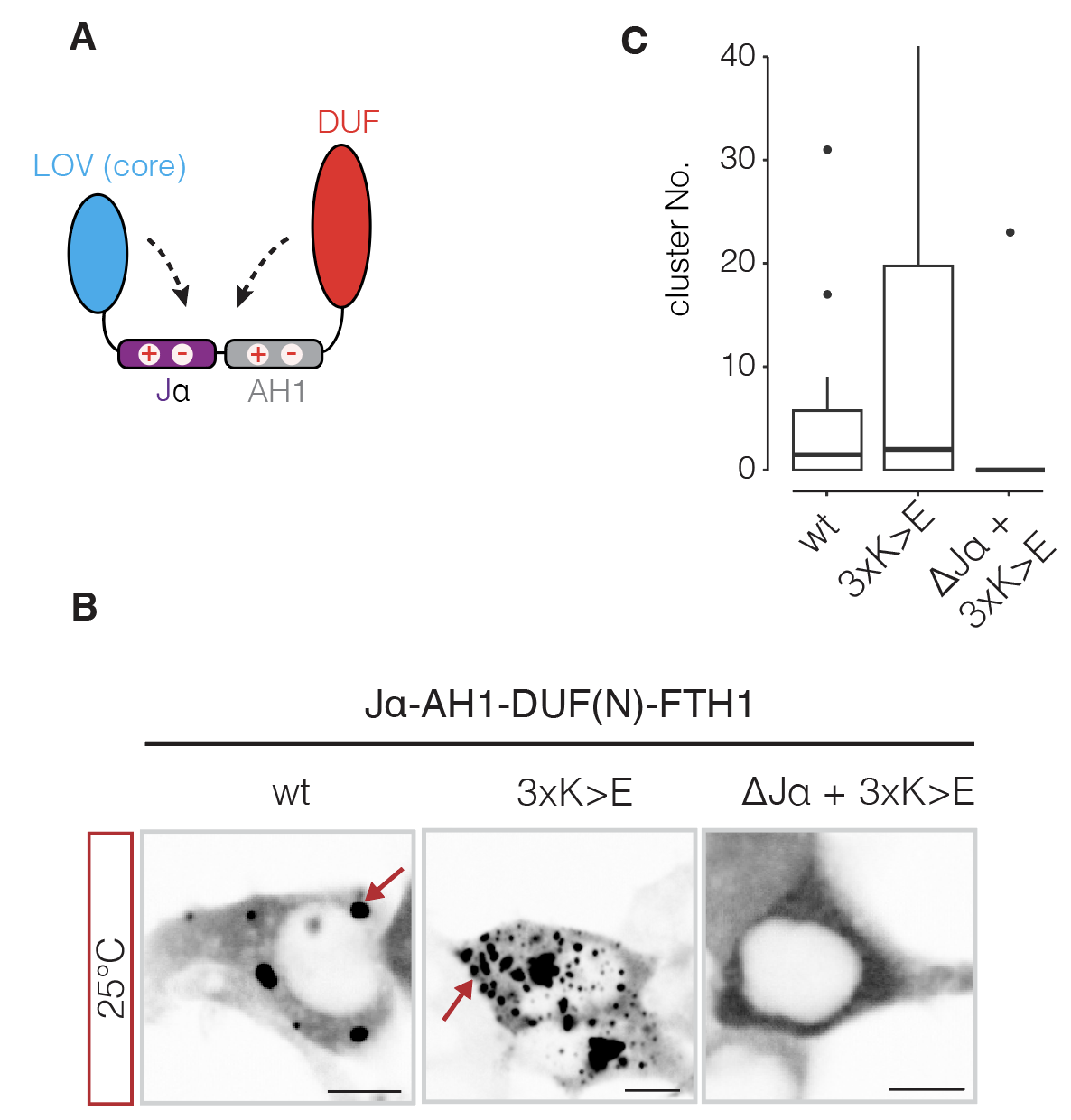

**Fig S9. Jα and balanced charge distribution are crucial for BcLOV4 clustering at 25°C.** (**A**) Jα-AH1-DUF(N) is the BcLOV4 fragment from amino acids 356 to 452, within which alternating positive and negative amino acid patches were found (**Figure 3N**). (**B**) HEK 293T cells were transfected with the indicated constructs and incubated at 25°C for one hour before imaging. 3K>E refers to introducing K to E mutations at K405, K410, and K414, which are important positions for membrane-binding that were identified previously. “∆Jα” refers to the wt fragment but excluding the Jα helix (a.a. 385-452). Scale bar = 10 µm. (**C**) Quantification of (**B**). 28-61 cells were quantified for each group.

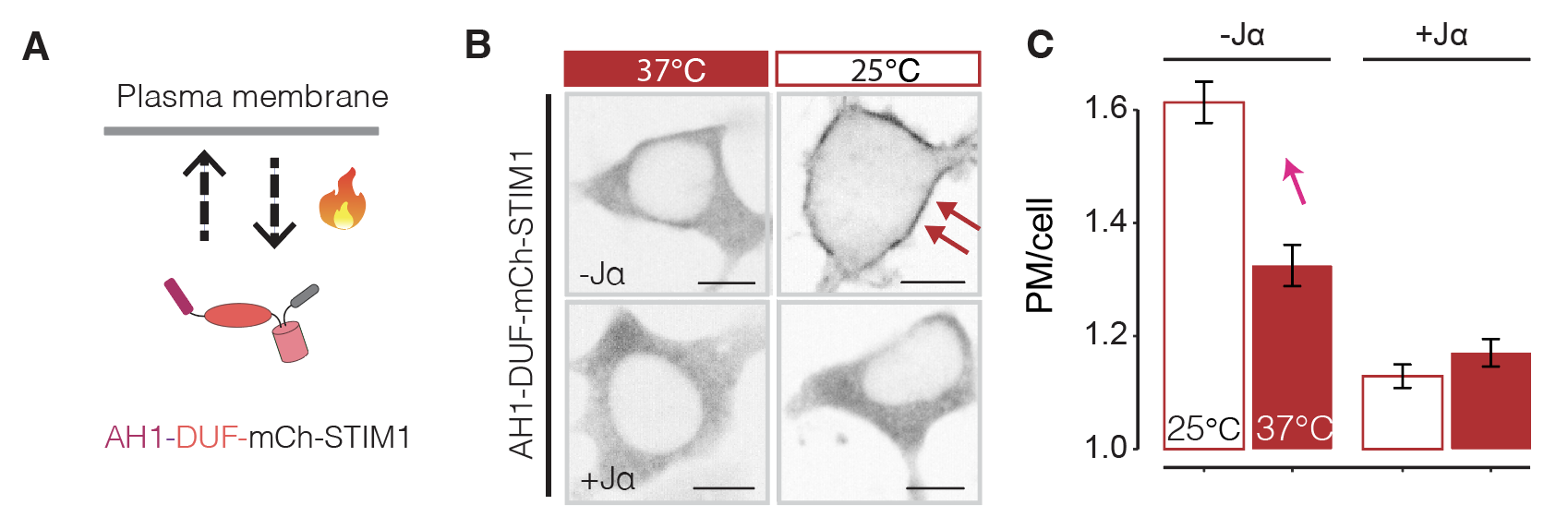

**Fig S10. DUF domain shows temperature sensitivity when fused to STIM1.** **(A)** AH1-DUF (385-595) was fused with STIM1 polybasic domain for testing temperature sensitivity. **(B)** HEK 293T cells were transfected with the indicated constructs with or without Jα (356-384) and imaged after pre-incubation at either 37°C or 25°C overnight. While the AH1-DUF fragment responds to temperature, addition of Jα suppresses switchability. Scale bar = 10 um. **(C)** Quantification of **(B)**. Data represent mean +/-SEM of ~300 cells.

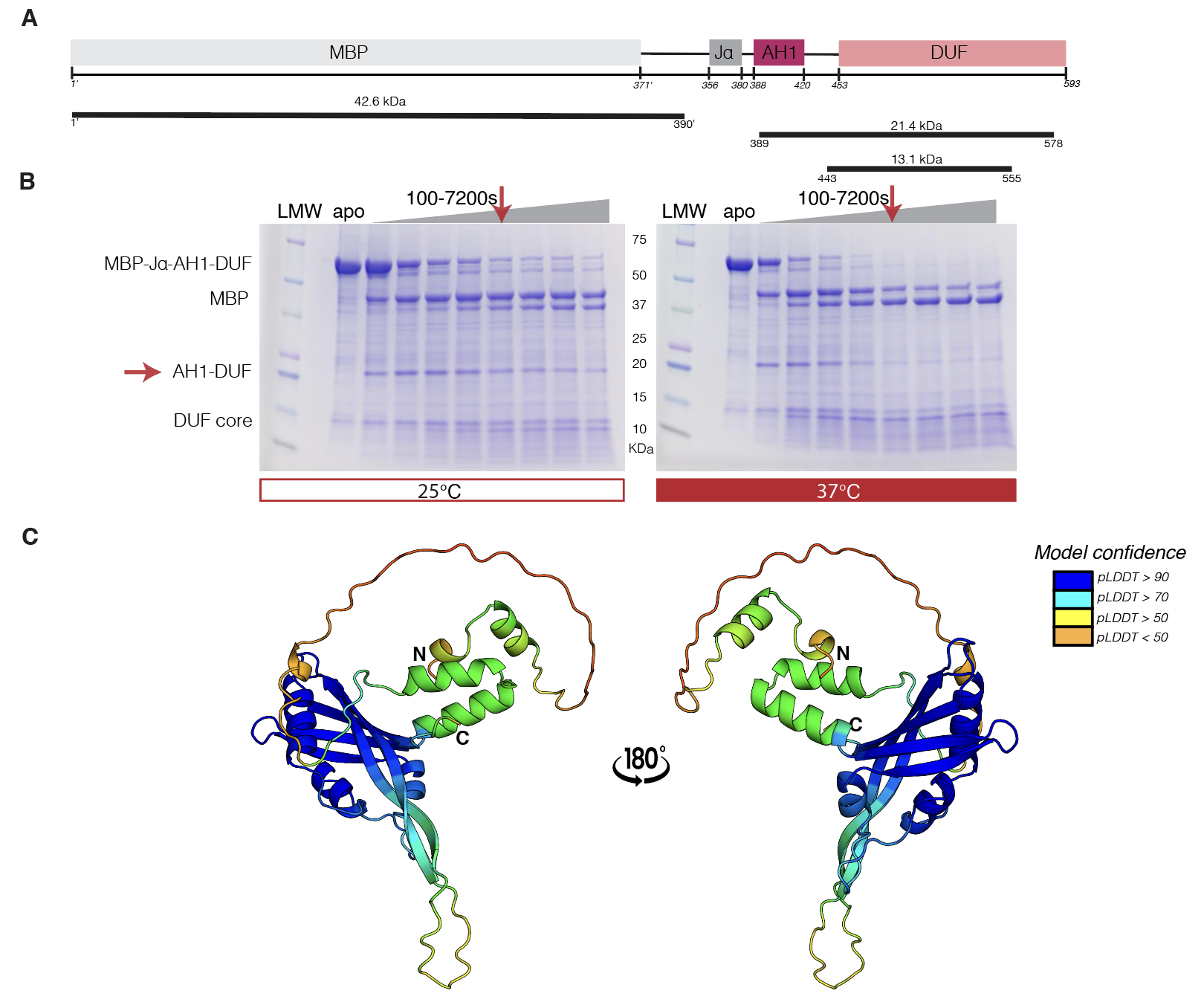

**Fig S11. Limited proteolysis reveals thermostability and thermoswitchability of the AH1-DUF region of BcLOV4.** (**A**) Domain schematic of MBP-Jα-AH1-DUF protein. Note that residues in the MBP and linker regions preceding Jα-AH1-DUF are designated differently (1’ - 403’) than those from Jα-AH1-DUF (numbered 356 - 593, as in full length BcLOV4). (**B**) SDS-PAGE gels show patterns of trypsin digestion of MBP- Jα-AH1-DUF at 25°C and 37°C over 7200 s. Cleavage of the soluble MBP tag from the MBP-Jα-AH1-DUF fusion is observed early under both conditions, with MBP staying thermostable under all conditions. AH1-DUF was moderately thermoresistant at 25°C, with some increased degradation at 37°C. A region predicted to be a thermostable part of the DUF domain was persistently observed at ~13.1 kDa. (**C**) AlphaFold3 model of Jα-AH1-DUF, colored by pLDDT confidence scores mapped onto structure.

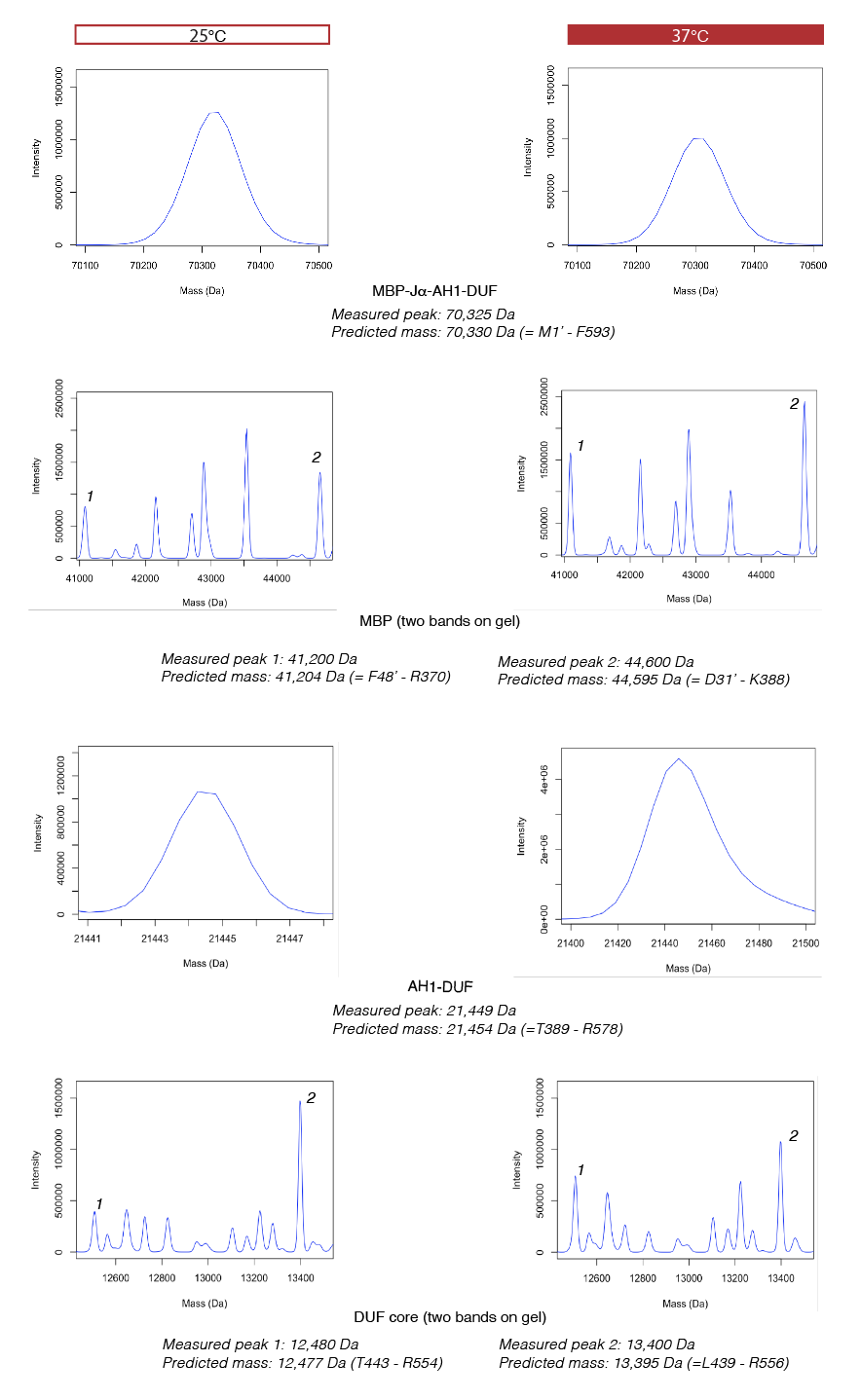

**Fig S12. Mass spectrometry data used for identifying fragments in limited proteolysis experiments of MBP-Jα-AH1-DUF.** Chromatograms depicting measured masses are shown for various bands that appear prominently in SDS PAGE gel. Note that residues in the MBP and linker regions preceding Jα-AH1-DUF are designated differently (1’ - 403’) than those from Jα-AH1-DUF (numbered 356 - 593, as in full length BcLOV4).

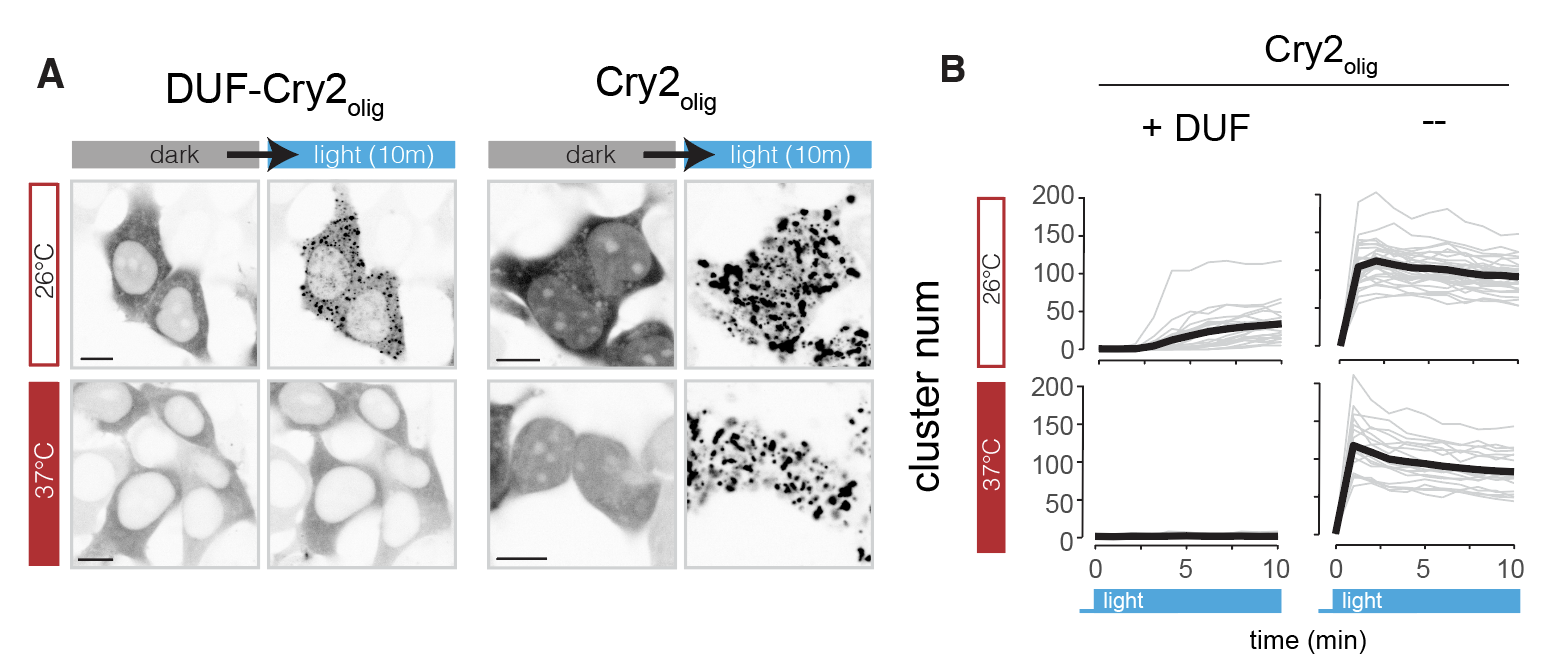

**Fig S13. Modular thermal regulation of unrelated proteins with DUF.** (**A**) HEK 293T cells were transfected with either DUF-Cry2_olig_ or Cry2_olig_. Cells were stimulated with blue light under 37°C or 26°C (2 hours pre-incubation). Scale bar = 10 µm. (**B**) Quantification of (**A**). Cluster number in each cell is quantified under each of the four conditions. Grey traces represent single cells, with black traces showing the mean. 20-30 cells were quantified for each condition. DUF-Cry2_olig_ data is reproduced from **Figure 4H,I.**

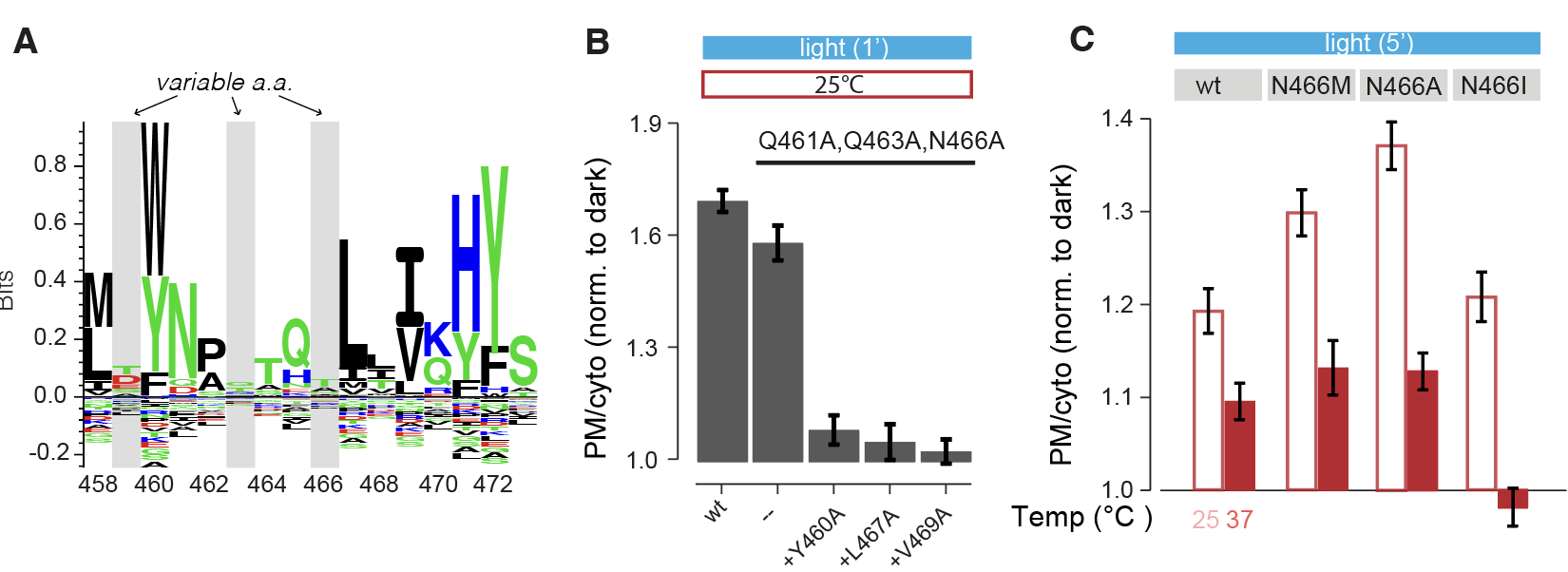

**Fig S14. Evolutionarily varied amino acids modulate BcLOV4 temperature sensing.** (**A**) Amino acid conservation of the BcLOV4(354-595) region among 69 homologues shows highly conserved and highly variable amino acids. (**B**) Mutations in conserved amino acids around position 465 resulted in poorly light responsive proteins. The degree of membrane binding after 1 minute of light stimulation was quantified. Whereas a triple mutation of non-conserved amino acids had little effect, additional mutation of any of three conserved amino acids substantially reduced light-dependent activation in HEK 293T cells. Cells were incubated under 25°C for 2 hours prior to stimulation. Data represent mean +/- SEM of ~150 cells. (**C**) N466 was replaced with amino acids found in BcLOV4 homologues (**Fig 4H**). Mutants were transfected into HEK 293T cells and were incubated under the specified temperatures for 2 hours prior to light stimulation. The identity of the amino acid mutation could either increase or decrease the magnitude of activation at high temperatures. Data represent mean +/- SEM of ~180 cells.

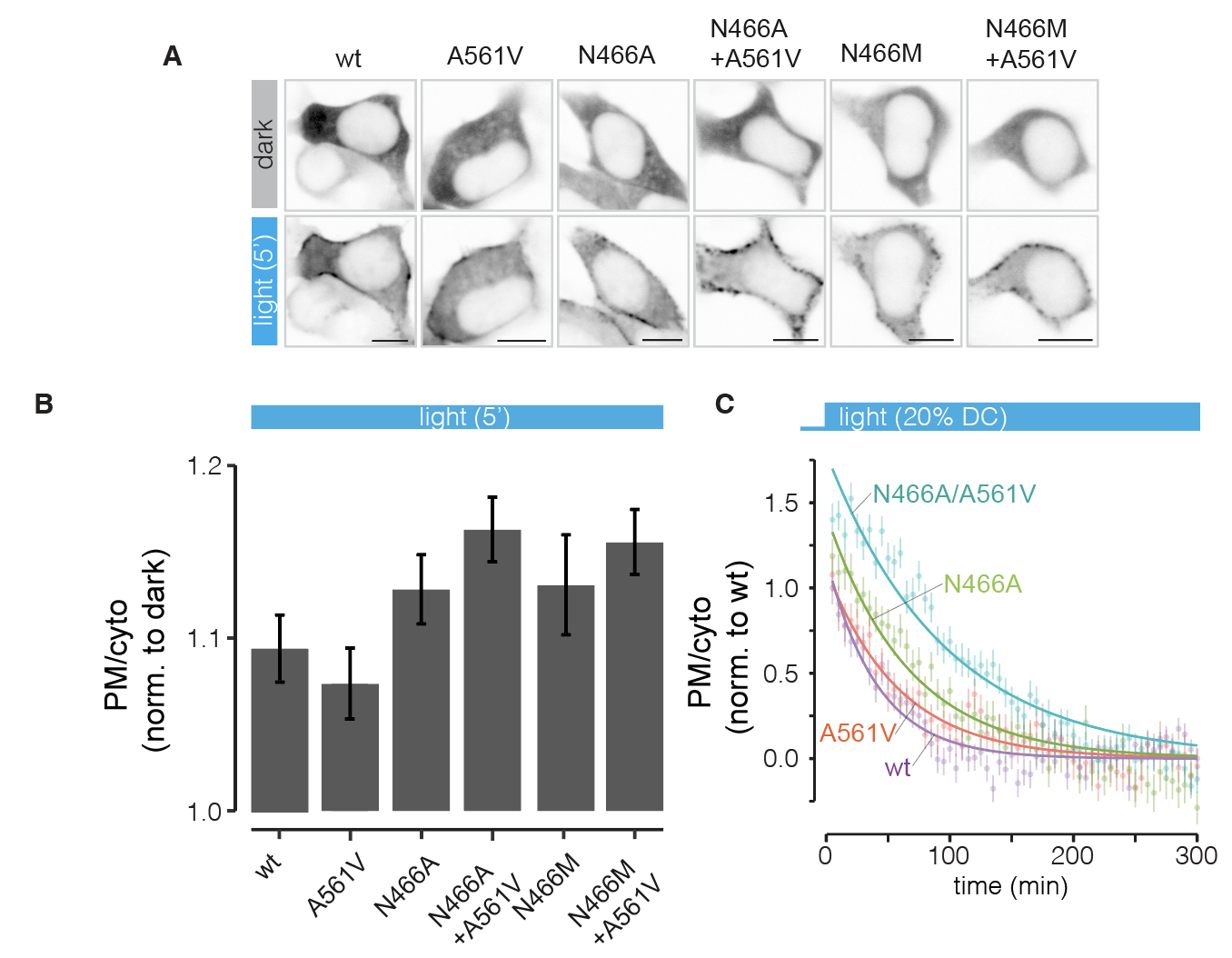

**Fig S15. Synergistic enhancement of membrane binding by addition of A561V to N466 variants.** Constructs were expressed in HEK 293T cells, which were incubated at 37°C after transfection. (**A**) Representative images of BcLOV4 variants light response. Scale bar = 10 µm. (**B**) Quantification of (**A**). Data represent mean +/- SEM of ~220 cells. (**C**) Time course of long-term light stimulation of BcLOV4 variants with replacement of co-evolved amino acids from homologs. Data are normalized to the max and min of membrane translocation for the wt BcLOV4 protein. Light was delivered through optoPlate with 20% duty cycle and 12 mW/cm^2^. Data represent mean +/- SEM of ~250 cells.

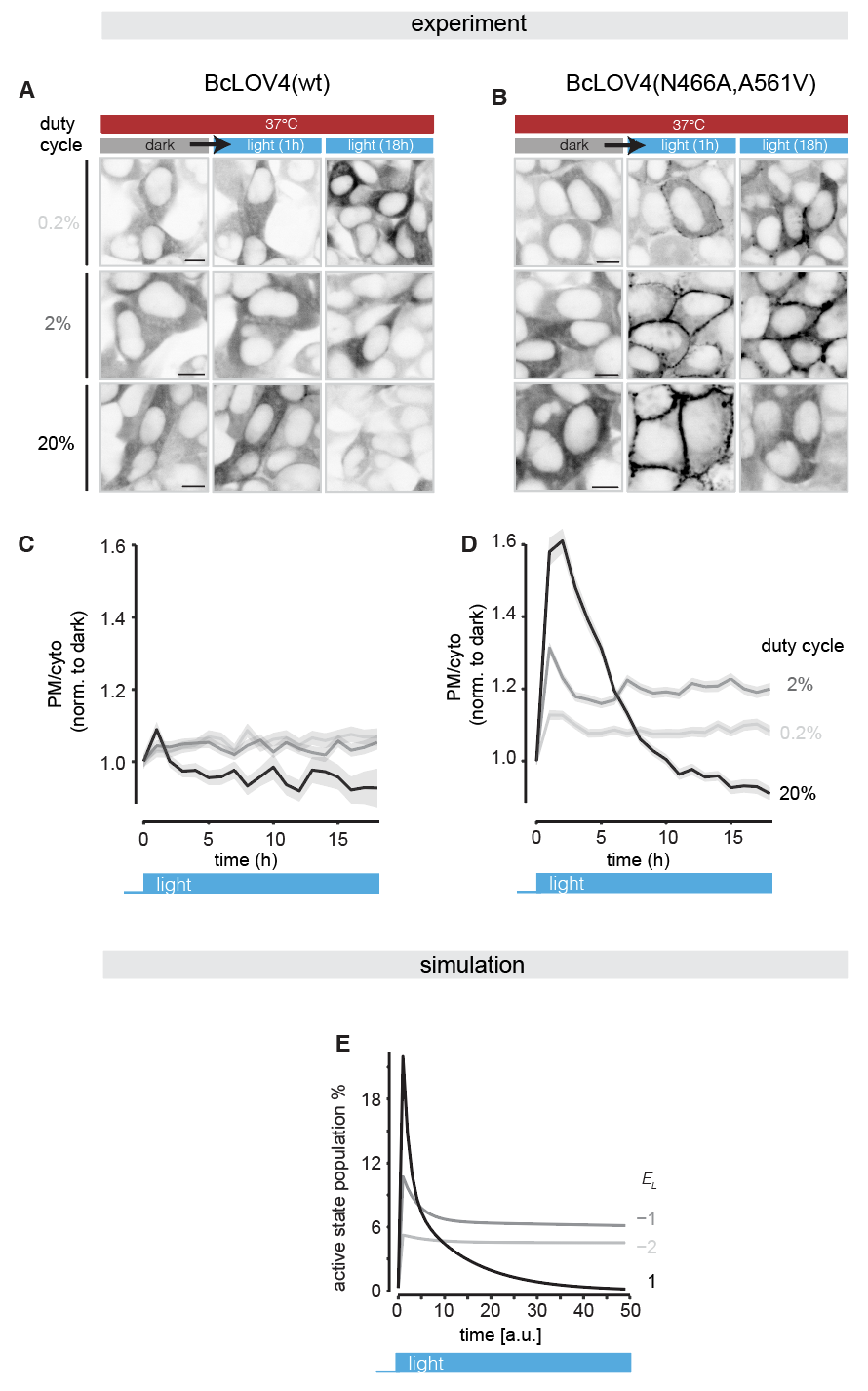

**Fig S16. Stable long-term activation of the N466A/A561V double mutant occurs under only low light stimulation.** (**A,B**) HEK 293T cells stably expressing either wt BcLVO4 or the N466A/A561V double mutant and were illuminated at variable duty cycles over 18 hours using the optoPlate (see Methods for details). Cells were cultured under 37°C. Light intensity stimulation intensity was 12 mW/cm^2^ and was applied at variable duty cycles. Scale bar = 10 µm. (**C,D**) Quantification of (**A,B**). The intersections between 20% duty cycle trace and others indicate a higher rate of entering irreversible state under higher duty cycles. Data indicate mean +/- SEM of ~ 280 cells. (**E**) The kinetic model (**Fig S1**) captures stable activation under intermediate stimulation but full decay under stronger stimulation, further supporting a light-dependent inactivation step.

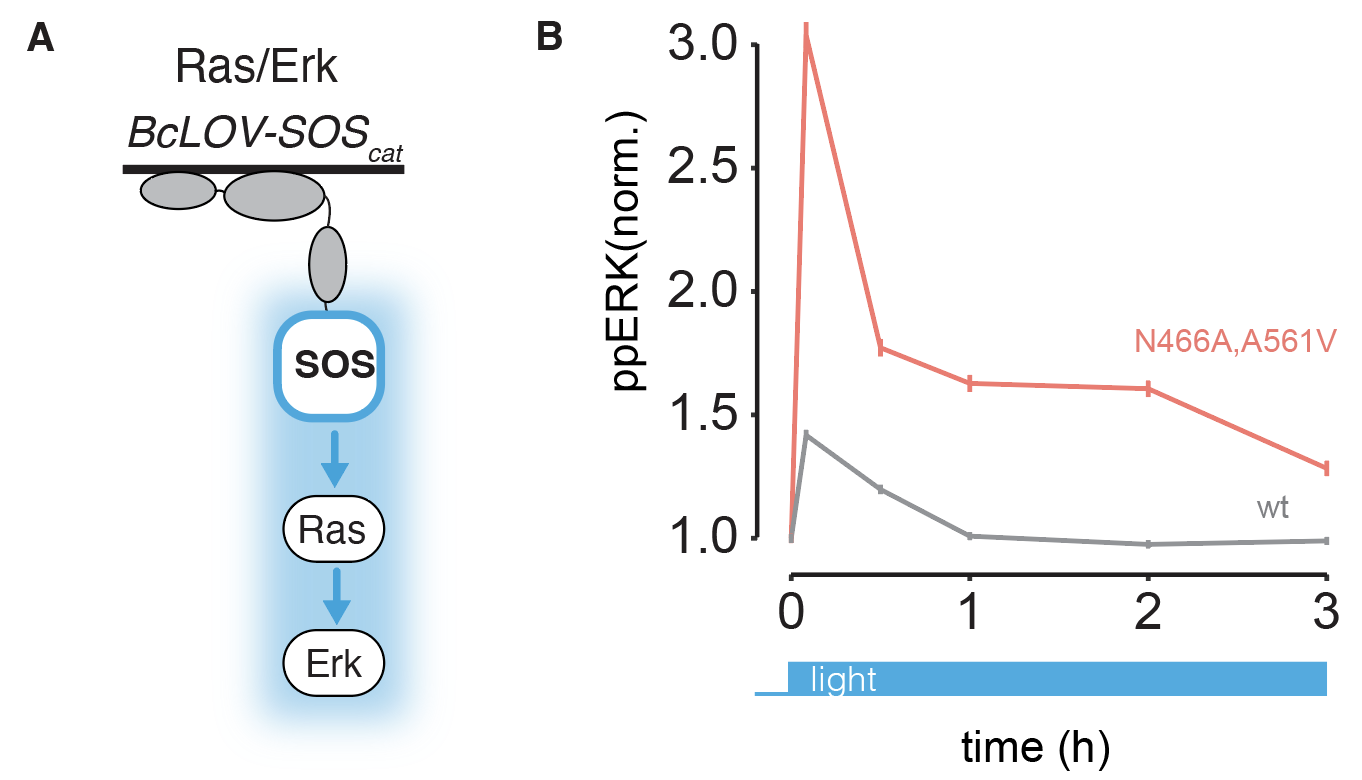

**Fig S17. DUF domain mutants enhance the potency and thermostability of BcLOV4-based signaling probes.** (**A**) Ras-Erk signaling can be controlled by translocation of the SOS catalytic domain to plasma membrane (BcLOV4-SOS_cat_). (**B**) NIH 3T3 cells stably expressing BcLOV4-mCh-SOS_cat_ (wt or N466A/A561V) were serum starved overnight and then stimulated with blue light (3.9 mW/cm^2^, 30% duty cycle) at 37°C. Data show immunofluorescence for phospho-ERK (ppERK) as a marker of pathway activity. Data represent mean +/- SEM of ~4500 cells per data point. The double mutant allows more potent and durable stimulation of the pathway relative to wt BcLOV4.

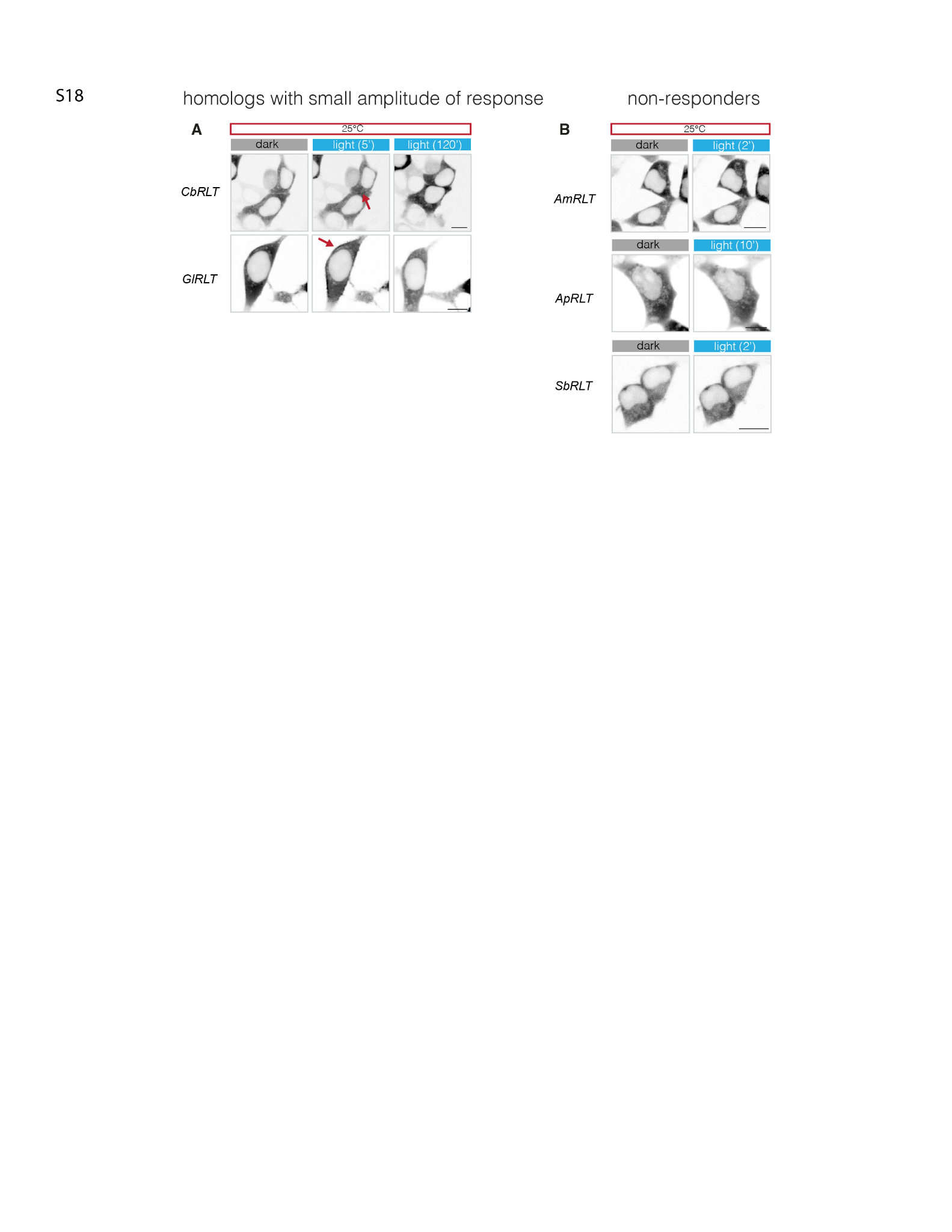

**Fig S18. Weak- or non-responders among BcLOV4 homologs.** Representative images of the rest of the synthesized homologues from **Figure 5G**. (**A**) The homologues with small response amplitudes under 25°C. Arrows highlight the membrane associated protein. (**B**) Homologues without obvious light response under 25°C. Scale bar, 10 um.

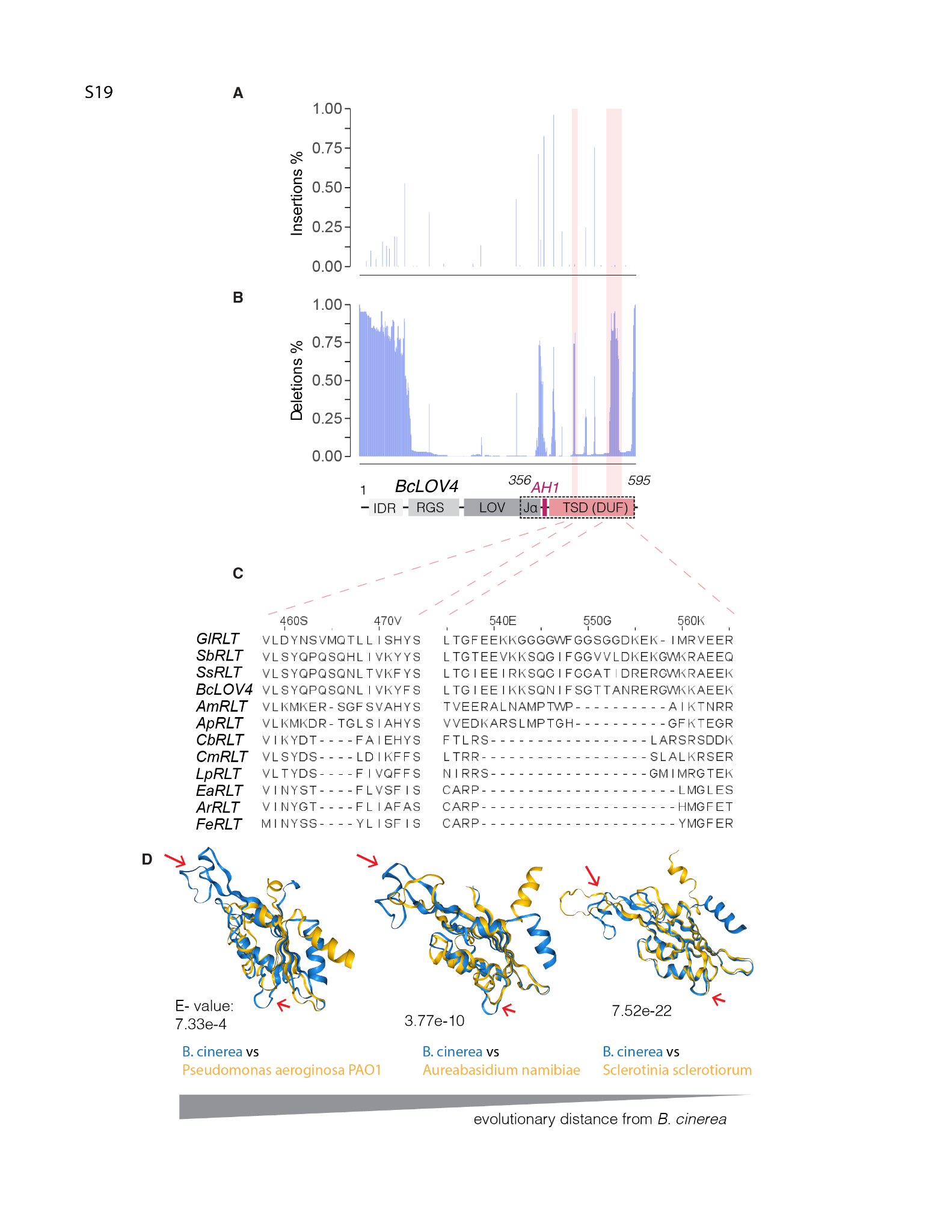

**Fig S19. Loop expansion in TSD over evolution.** (**A**) Sequences from 230 evolutionarily related BcLOV4 homologues were analyzed for insertion events, using BcLOV4 as the reference. (**B**) The same list of sequences in (**A**) was used to detect deletion events using BcLOV4 sequence as reference. Peaks identified in both (**A**) and (**B**) indicate mismatches between homologues and BcLOV4. Regions with only deletion peaks were highlighted, indicating evolutionary sequence expansion events during BcLOV4 evolution. This analysis excludes deletions at the beginning and end of the reference, as these occur because of a lack of surrounding aligned sequence for calling insertions. Note: insertion peaks appear narrower than deletion peaks because insertions specify the amino acid position where insertions were made, whereas deletions can describe a large region of deleted residues. (**C**) The detailed sequence of expanded regions in BcLOV4 and homologues synthesized for testing. (**D**) Aligned folds of TSD from BcLOV4 (blue) and homologues (yellow) searched with Foldseek. The E-value was annotated for indicating evolutionary distance (smaller E-value indicates a more recent divergence). Red arrows highlight the loop regions expanded along evolution.

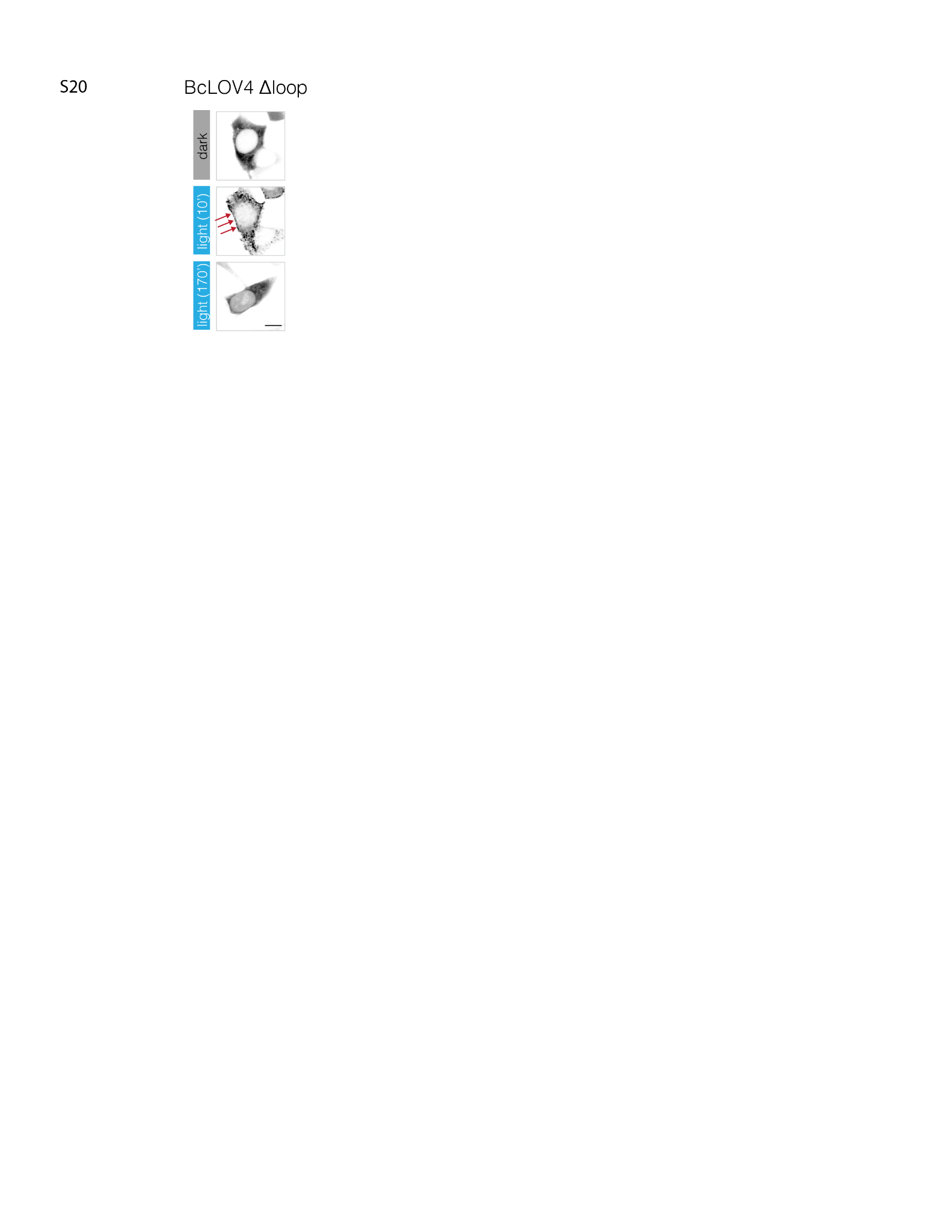

**Fig 20. BcLOV4 Δloop responds to light with pulsatory activity at a shifted temperature range.** The light and temperature condition is indicated by the white circle in **Figure 5H**. Scale bar, 10 um.

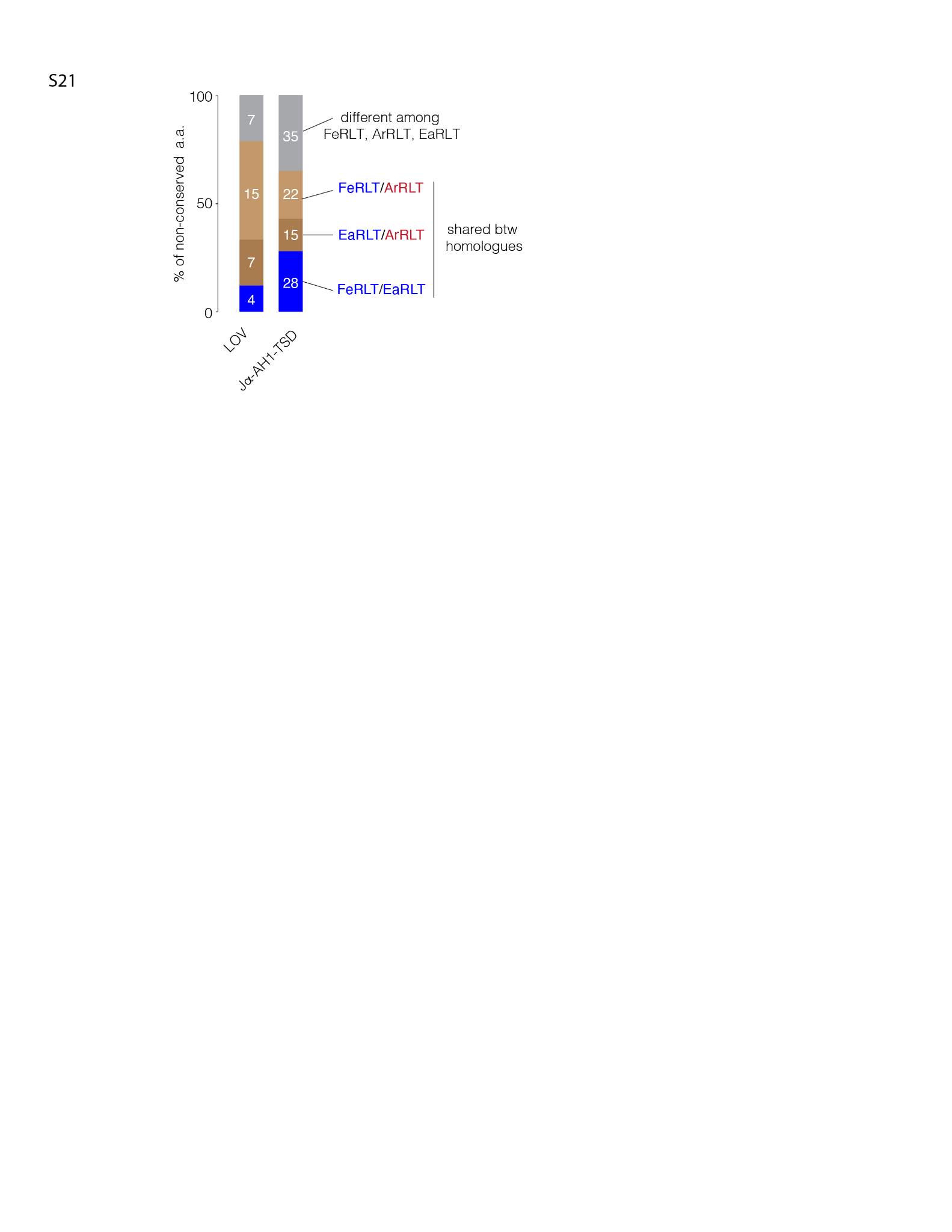

**Fig S21. Convergent evolution of TSD during cold adaptation.** The LOV domain and the Jα-AH1-TSD region of the homologues were used to quantify similarities in the non-conserved amino acids (among FeRLT, EaRLT, ArRLT) among homologues. Cold-adapted homologues are colored in blue while hot-adapted is in red. Although FeRLT and ArRLT diverged most recently in evolution, the cold-adapted homologues FeRLT and EaRLT share more of the non-conserved amino acids in Jα-AH1-TSD, suggesting their convergent adaptation to cold environments. The number in the box indicates the absolute number of amino acids counted in each group.
